# Genetic dissection of root-mediated yield heterosis in melon (*Cucumis melo*)

**DOI:** 10.64898/2026.04.15.718623

**Authors:** Asaf Dafna, Galil Tzuri, Elad Oren, Tal Isaacson, Ilan Halperin, Gadi Peleg, Amit Gur

## Abstract

Heterosis, the superiority of hybrids over their parents, is a major genetic force associated with plant fitness and crop yield enhancement. We previously discovered and characterized root-mediated yield heterosis (RMYH) in melon (*Cucumis melo*) using a half-diallel population, derived from 20 diverse parents. In the current study we investigated the genetic architecture of RMYH using a segregating population derived from a selected F_1_ hybrid (HDA019) that consistently induced RMYH under several melon scion varieties and growing conditions. 78 recombinant inbred lines (RILs) and their test-crosses to both parents were analyzed in yield trials as rootstocks under a common commercial scion variety. The population displayed normal root-mediated yield distribution and transgressive segregation relative to the parents but none of the RILs equaled the superior performance of the F_1_ hybrid. RMYH of HDA019 was dissected to small effect QTLs showing mostly additive or dominant mode-of-inheritance and favorable QTL-alleles were contributed by both parents. Five consistent QTLs were selected and used to demonstrate the potential of root-mediated yield QTL pyramiding, and 20 combinations of QTL pairs and triplets supported the cumulative model for heterosis. Favorable QTLs alleles were introgressed to generate advanced QTL-backcross lines that were used for validation. This study provides first detailed genetic dissection of yield-related rootstock traits in cucurbits, highlighting rootstock breeding as an important underutilized route for improving yield and stress tolerance of crops.

**Key message:** Root–mediated yield heterosis in melon was genetically dissected using grafting strategy, revealing additive QTLs from both parents of the mapping population. Rootstock breeding through pyramiding of favorable alleles is proposed as strategy for enhancing crop yield and stress tolerance.

## Introduction

Global food security requires continued increases in crop productivity while reducing environmental cost, which motivates new routes to improve yield stability and resource□use efficiency (Godfray *et al*., 2010; Tilman *et al*., 2011; FAO, 2025). Conventional breeding has primarily targeted above□ground traits, but further gains are likely to depend also on underexploited sources of variation such as root system function, which determines water and nutrient capture and mediates plant responses to stress (Parry and Hawkesford 2010; Lynch 2019). Root□targeted approaches therefore offer a complementary breeding strategy to increase yield resilience without altering desirable scion quality traits (Lynch 2019).

Roots control multiple processes that limit crop performance—water and mineral uptake, synthesis and transport of growth regulators, and interactions with soil microbiota—and therefore shape whole⍰plant responses to abiotic stress and spatial and temporal variation in soil resources (Fitter 2002; Hodge 2004; Smith and De Smet 2012). Root system architecture (RSA), the spatial configuration of the root system in soil, is highly plastic and can determine resource⍰capture efficiency and productivity under stress; variation in RSA has been linked to yield differences across crops (Zhu et al. 2005; Smith and De Smet 2012; Uga et al. 2013). Despite their importance, root traits remain underexploited in breeding programs due to multiple challenges. Field phenotyping of root systems is technically demanding and labor-intensive, while the genetic architecture underlying root traits is typically polygenic with substantial genotype × environment (G×E) interactions that complicate selection. Moreover, the complexity of root-shoot communication, mediated through hydraulic signals, hormonal transport (including abscisic acid, cytokinins, and strigolactones), and electrical/Ca²⁺ signaling, makes it difficult to predict how root modifications will affect above-ground performance (Sakakibara 2006; Gomez-Roldan et al. 2008; Osakabe et al. 2014; Huntenburg et al. 2022). Ongoing advances in non⍰invasive phenotyping, genomics and experimental designs (including grafting and field coring/portable imaging) now enable field⍰relevant assessment of root function using scion agronomic endpoints (yield, quality), making selection on root⍰expressed effects more feasible (Wasson et al. 2014; Lynch 2019; Tracy et al. 2020).

Heterosis, or hybrid vigor, denotes the phenomenon in which F_1_ hybrids outperform their parents in one or more traits (East 1908; Shull 1908). Since its systematic adoption in hybrid maize breeding early in the 20th century, heterosis has been central to crop improvement and underpins a large fraction of modern yield gains (Duvick 2001). The genetic basis remains unresolved—dominance complementation, overdominance and epistasis have all been proposed—and recent genomic and QTL analyses emphasize polygenic architectures with many sma⍰lleffect loci and important epistatic interactions (Semel et al. 2006; Lippman and Zamir 2007; Schnable and Springer 2013; Huang et al. 2015; Liu et al. 2020; Xiao et al. 2021). While heterosis has been studied and exploited primarily for above⍰ground traits, its potential expression in root systems, root⍰mediated heterosis, remains comparatively unexplored and represents a promising, under⍰used avenue to improve whole⍰plant performance.

Understanding the genetic basis of root traits is essential for their effective manipulation in breeding programs. Studies in model plants and major crops show that root phenotypes are generally quantitatively inherited and highly environment⍰sensitive (Smith and De Smet 2012). QTL mapping and meta⍰analyses have revealed abundant heritable variation; for example, hundreds of root⍰ and stress⍰related QTLs compiled in rice meta⍰analyses (Courtois et al. 2009; Satasiya et al. 2024) and numerous maize QTLs for lateral rooting, root⍰hair and other architectural traits (Zhu et al. 2005). However, most mapped loci have sma⍰llto⍰moderate effect and display G×E interactions. Despite this complexity, mapping has yielded identifiable genes with clear agronomic effects—for example, *DRO1*, which controls root angle and improves drought avoidance (Uga et al. 2013), and *PSTOL1*, which enhances early root growth and phosphorus acquisition (Gamuyao et al. 2012). Advanced genetic mapping populations and genomic platforms increasingly enable high⍰resolution trait dissection and candidate⍰gene discovery (McMullen et al. 2009; Huang et al. 2015), but progress requires integrated field⍰relevant phenotyping and multi⍰environment validation (Zhu et al. 2005; Lynch 2019). Consequently, crops and systems that permit field⍰scale root–shoot dissection are particularly valuable for translating allelic variation into agronomic gains.

Melon, *Cucumis melo* L. (Cucurbitaceae) is an economically important horticultural crop with global production averaging approximately 28.2 million tons per year (mean 2021–2024; FAO, 2025). The species exhibits extensive intraspecific diversity: two major subspecies (ssp. *melo* and ssp. *agrestis*) and many botanical varieties and market classes provide abundant allelic variation for breeding (Pitrat 2017; Gonzalo et al. 2019). *Cucumis melo* is a diploid with 12 chromosomes (n=12) and an estimated genome size of ∼400-450 Mbp (Garcia-mas et al. 2012). In addition to the extensive genetic resources already available in melon (*e.g.* diverse collections and segregating populations), the sequencing of the melon genome, completed in 2012 (Garcia-Mas et al. 2012), established a robust foundation for advanced genomic investigations. This milestone has facilitated whole-genome resequencing efforts encompassing more than 1,000 diverse melon accessions (Zhao et al. 2019) for detailed description of genetic variation within this species. Recently, additional melon genomes were *de novo* assembled, and a pan-genome framework is now available for more detailed comparative genomic analyses (Oren et al. 2022). Among other horticulturally-relevant traits, root traits in melon are amenable to genetic analysis. Near⍰isogenic and mapping studies document heritable variation in lateral branching, root⍰hair and other architectural components (Fita et al. 2008), providing opportunities to identify alleles that influence below⍰ground variation and their potential effect on above-ground traits.

Grafting is widely used in cucurbit production to control soi⍰lborne diseases and improve stress tolerance and is simple in melon, enabling the experimental separation of rootstock and scion genotypes in the field (Flores-León et al. 2021). Importantly, rootstocks can alter scion vegetative vigor, yield and fruit quality through root⍰mediated effects (Alarcón et al. 2020), and the availability of diverse collections and cultivars facilitates identification and deployment of beneficial underutilized root alleles (Gonzalo et al. 2019). These properties make melon grafting an attractive, translational route to move root-related genetic variation into agronomic gains while retaining scion traits.

In a previous work we demonstrated root⍰mediated yield heterosis in melon: a survey of hybrid rootstocks revealed best⍰parent heterosis (BPH) for scion yield ranging from 25% to 79% (Dafna et al. 2021). A highly heterotic F_1_ rootstock (HDA019; the hybrid of PI414723 × DUL) consistently increased scion yield relative to both parental rootstocks. Building on this finding, in this study we dissected the genetic architecture underlying root⍰mediated yield heterosis of HDA019 by exploiting a PI414 × DUL RILs population and derived reciprocal test⍰cross populations. We mapped and prioritized consistent root-mediated yield (RMY) QTLs, evaluated QTL interactions, and validated selected QTLs by development and testing of advanced QTL backcross lines. The results indicate predominantly additive and dominant contributions and demonstrate that QTL stacking can reproduce a fraction of the F_1_ hybrid rootstock advantage, supporting hybrid rootstock breeding as a practical route to improve yield performance, while retaining the varietal properties of the scion.

## Materials and Methods

### The germplasm, population development and experimental designs are illustrated in ***Figure 1***

### Plant material

### Bi-parental recombinant inbred lines (RILs) and test-cross (TC-RILs) populations

A RIL population (PI414 × DUL, F_6-8_), developed from a cross between ‘Dulce’ (DUL; *C. melo* subsp. *melo* var. *reticulatus*) and ‘PI414723’ (PI414; *C. melo* subsp. *agrestis* var. *momordica*) was previously developed (Danin-Poleg et al. 2002; Harel-Beja et al. 2010) and used in this work. The population was genotyped with 58,328 genome-wide SNPs (Galpaz et al. 2018); 78 RILs with low heterozygosity levels were selected for this study. Two test-cross (TC-RILs) populations were built. Plants of the 78 RILs were grown and crossed to the parents DUL and PI414 in the greenhouse at Newe-Ya’ar Research Center, Israel (32°43’05.4”N 35°10′47.7″E) during the fall of 2018.

**Figure 1:**
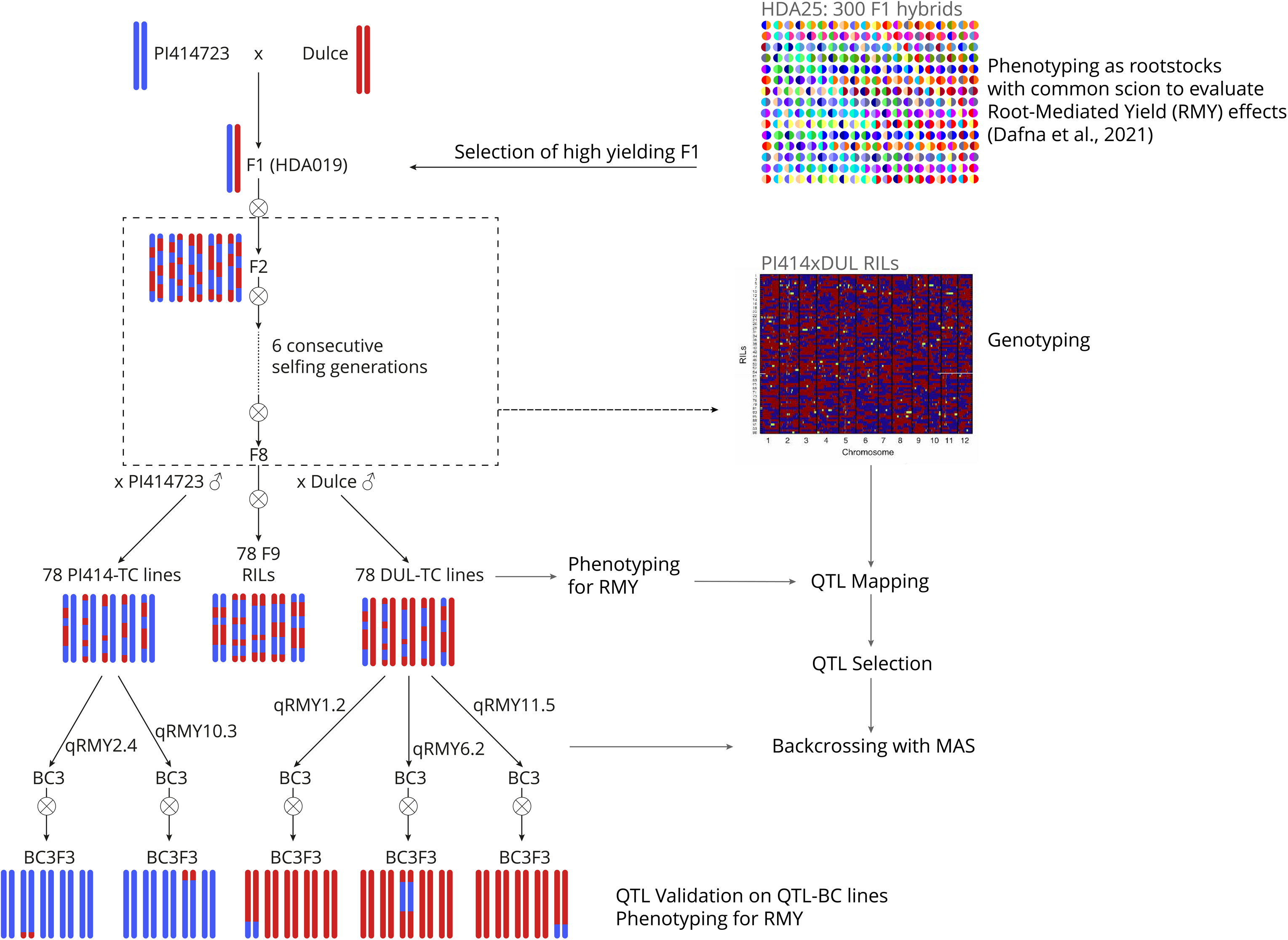
Schematic outline of the experimental workflow and germplasm development. Development of the PI414 × DUL RIL, TC-RILs and QTL-BC lines populations for root-mediated yield (RMY) analyses. The F_1_ hybrid (HDA019; high-yielding F_1_ rootstock) was generated by crossing PI414723 (blue) and Dulce (red). An F8 recombinant inbred line (RIL) population was developed through consecutive generations of selfing (Danin-Poleg et al. 2002; Harel-Beja et al. 2010). The RILs were then test-crossed to both parental lines to create two TC-RIL populations (78 PI414-TC lines and 78 DUL-TC lines) for phenotyping. The RIL populations was genotyped with genome-wide SNPs (Galpaz et al. 2018). The RIL and TC-RIL populations were phenotyped for RMY in a replicated grafted experiments over two field season. QTL mapping was performed to identify loci associated with RMY variation. Selected QTLs were introgressed into the parental backgrounds through marker-assisted backcrossing, generating QTL-BC lines for each QTL. These QTL-BC lines were further phenotyped to validate the effects of the introgressed QTLs. MAS: Marker-assisted selection.

### Creation and testing of root-mediated yield QTL-BC lines

The objective was to create advanced QTL backcross (QTL-BC) lines of each of 5 selected consistent RMY QTLs through introgression of their QTL intervals from the favorable allele donor parent to the background of the recurrent parent.

Ten lines, out of the 78 RILs, representing 2 lines for each QTL, were selected for marker-assisted backcrossing. Criteria for selection of these lines was based on the presence of favorable allele in the complete QTL-confidence interval, defined by decrease of 2-LOD scores flanking the QTL peak, and the highest proportion of the recurrent parent alleles in the rest of the genome, based on the aforementioned genome-wide genotyping of the RILs population.

These lines were grown and crossed to their respective recurrent parent in the greenhouse at Newe-Ya’ar Research Center, Israel. Starting with the F_1_ of the TC-RILs, we performed three consecutive backcrossing to the recurrent parent in the spring (BC_1_) and summer (BC_2_) of 2021 and spring (BC_3_) of 2022. The selected BC_3_ seeds contain approximately 94% recurrent parent genome. Prior to each backcrossing cycle from the BC_1_ and onwards, for each QTL-BC line 92 seeds were genotyped (see genotyping section) and only seeds that were heterozygous throughout the complete QTL confidence interval (approx. 10-15 seeds) were selected for sowing. In the summer of 2022, we selfed the BC_3_ lines to produce BC_3_F_2_ seeds.

For the field trials of the advanced QTL-BC lines, we tested plants homozygous for the favorable or non-favorable alleles in the introgression as well as heterozygous, and therefore BCF_2_ seeds were genotyped pre-sowing and were then grown and used as rootstocks for the grafted plants. The selected seeds were sown in a commercial nursery and grafted with ‘Glory’ scion using standard nursery protocols. However, due to germination and grafting issues, fewer than 300 grafted seedlings were obtained, with 12-25 seedlings representing most genotypes. The grafted plants were planted in the open-field at Newe-Ya’ar, using a randomized block design. Parental and F_1_ rootstocks, as well as non-grafted ‘Glory’ plants, served as controls.

### Rootstock grafted yield trials

The results described in the current manuscript were collected over 19 field experiments that were conducted from 2017 to 2024 (**Supplementary Table S1**). Each genotype (from either the RILs, TC-RILs, QTL Back-cross lines, parental lines or controls) was grafted as a rootstock with a common scion. Grafting for these large-scale experiments was performed in a commercial nursery (’Hishtil’—Ashkelon, Israel, https://www.hishtil.com/) under their standard grafting protocols and the field trial was carried out as described in (Dafna et al. 2021). Briefly, Rootstocks and scions were separately sown, grafted after 21 days, and ready for field transplanting in 7-10 days. The common scion used was ‘Glory’, a ‘Galia’-type variety with uniform fruit setting. For field trials, plants were grown in a randomized complete block design, harvested once when >70% of the fruits were fully ripe, and 95% reached their full size. Replicated trials consisted of five plots of five plants for each genotype. Average fruit weight (AFW) and total soluble solids (TSS) were measured for yield and quality analysis. In each plot, all fruits were harvested, counted and weighed, and Total Yield was assessed as the weight of the full size fruits per plant or plot. Average fruit weight was calculated by dividing the total fruit weight by the total number of fruits (FN) per plot.

### Root development and biomass analysis of DUL, PI414 and their F_1_ (HDA019)

Root phenotyping was performed by PhenoRoot Ltd. (Rehovot, Israel, https://www.phenoroot.com/). The hybrid HDA019 and its parental lines (Dulce, PI414723) were grown in pots under greenhouse conditions (n = 4 plants per genotype per time point). Destructive sampling was conducted at 7 days intervals across four time points. At each sampling, roots were separated from the soil by shaking on a mesh screen, washed, blotted dry, and weighed to obtain fresh root biomass. Root biomass distribution by depth was determined by partitioning roots into three segments (0–4, 4–8, and 8–12 cm depth) and weighing each fraction separately.

### Genotyping of QTL-BC lines

To confirm the genotypes of selected QTL-BC lines, fifteen InDel markers, in size range of 14-130 bp, that show polymorphism between the parents DUL and PI414 were detected and used as simple PCR markers as described in (Oren et al. 2022). Three markers per QTL were developed, focusing on the peak and both ends of the introgression. List of all the markers and primers is provided in **Supplementary Table S2**. Primers were designed using Gene Runner version 6.5. Genomic DNA was extracted from melon seeds in a non-destructive manner for pre-planting genotypic selection. From each seed, a small chip from the distal end (embryonic cotyledons side) of the seed was chopped off using a nail clipper. The small chips and the chopped seeds were placed in a parallel order in separate 96-well plates. The chopped-seed plates were wrapped with saran and kept in a 4°C refrigerator for as long as three months prior to sowing. DNA was extracted from the seed chips using a protocol modified from (Wang et al. 1993). Briefly, 75 µl of buffer A (100 mM NaOH + 2% Tween 20) were added to each well (without grinding), and the plate was then incubated for 10 minutes at 95°C. Then 75 µl of buffer B (100 mM Tris-HCl + 2 mM EDTA) were added, and the plate was agitated moderately for five minutes. Afterwards, 1–2 µl of the solution were used for PCR with the 2XPCRBIO HS Taq Mix Red (PCRBIOSYSTEMS, UK). The annealing temperature was 56°C. The products were separated on a 2.5% or 1.2% agarose gel for 1-2 hours. Based on the genotypic results, chopped seeds were selected from the corresponding plate positions and sown, subsequently germinating into normal-appearing plants, and displayed normal germination rates.

### QTL mapping and statistical analyses

JMP ver. 17.0.0 statistical package (SAS Institute, Cary, NC, USA) was used for statistical analyses. Mean yield comparisons were performed using the least square means (Entry mean) based on a fitted model that accounted for the effects of genotype, replicate and relative position in the field, including interactions between these factors. Broad-sense heritability (*H^2^* = (Var (G)/Var (P)) was estimated in each experiment separately using analysis of variance (ANOVA) based variance components. QTL analysis was performed as previously described in (Oren et al. 2020) In brief, TASSEL v.5.2.43 (Bradbury et al. 2007) was used for genome-wide single-marker analysis of the traits using generalized linear model (GLM). Interval mapping was performed with R/qtl (Broman et al. 2003). FDR method was used to control for multiple comparisons in linkage mapping. QTL effects were calculated as the yield difference between all lines carrying the favorable alleles and all lines with the non-favorable alleles based on an entry mean bases. Significance of the effects was determined by comparison of means. QTL interactions were analyzed in a two-way ANOVA test.

## Results

### HDA019 F_1_ displays root-mediated BPH and used for construction of mapping populations

In a recent study we used the diverse collection of melon accessions, which was built in the Cucurbits Unit at Newe Ya‘ar, to develop a half-diallel population, through structured intercrossing of core subset of 25 lines from this collection in all possible combinations (*HDA25*, (Dafna et al. 2021). The *HDA25* population served as a foundational resource to explore root function variation and its impact on whole-plant performance in field conditions, using a common scion grafting strategy, and to investigate the genetics underlying root-mediated yield variation (Dafna et al. 2021).

From the half-diallel study, we identified ‘HDA019’, an F_1_ hybrid between ‘Dulce’ (DUL; *Cucumis melo* ssp. *melo* var. Reticulatus) and PI414723 (PI414; C. *melo* ssp. *agrestis* var. Momordica), as one of the highest-yielding rootstocks. ‘HDA019’ consistently exhibited significant best-parent heterosis (BPH) for root-mediated yield (RMY), ranging from 25-79% across multiple grafted-rootstock yield experiments conducted over several years, under different growing conditions (Error! Reference source not found.**A, B; Sup. Table 1**).

In line with the consistent root-mediated yield heterosis of HDA019, in a root system architecture analysis experiment we found that root biomass and biomass accumulation rate of HDA019 were significantly higher than that of both DUL and PI414, with more than 100% BPH for this trait in different time points along the growing period (**Figure 2C**). Segmented by depth, HDA019 exhibited an intermediate root biomass distribution pattern compared to its parents, with a more balanced allocation across depths than Dulce, which concentrates roots in the upper layer, and PI414723, which shows greater tendency for deeper root development (**Figure 2D and Sup. Figure 1**).

**Figure 2:**
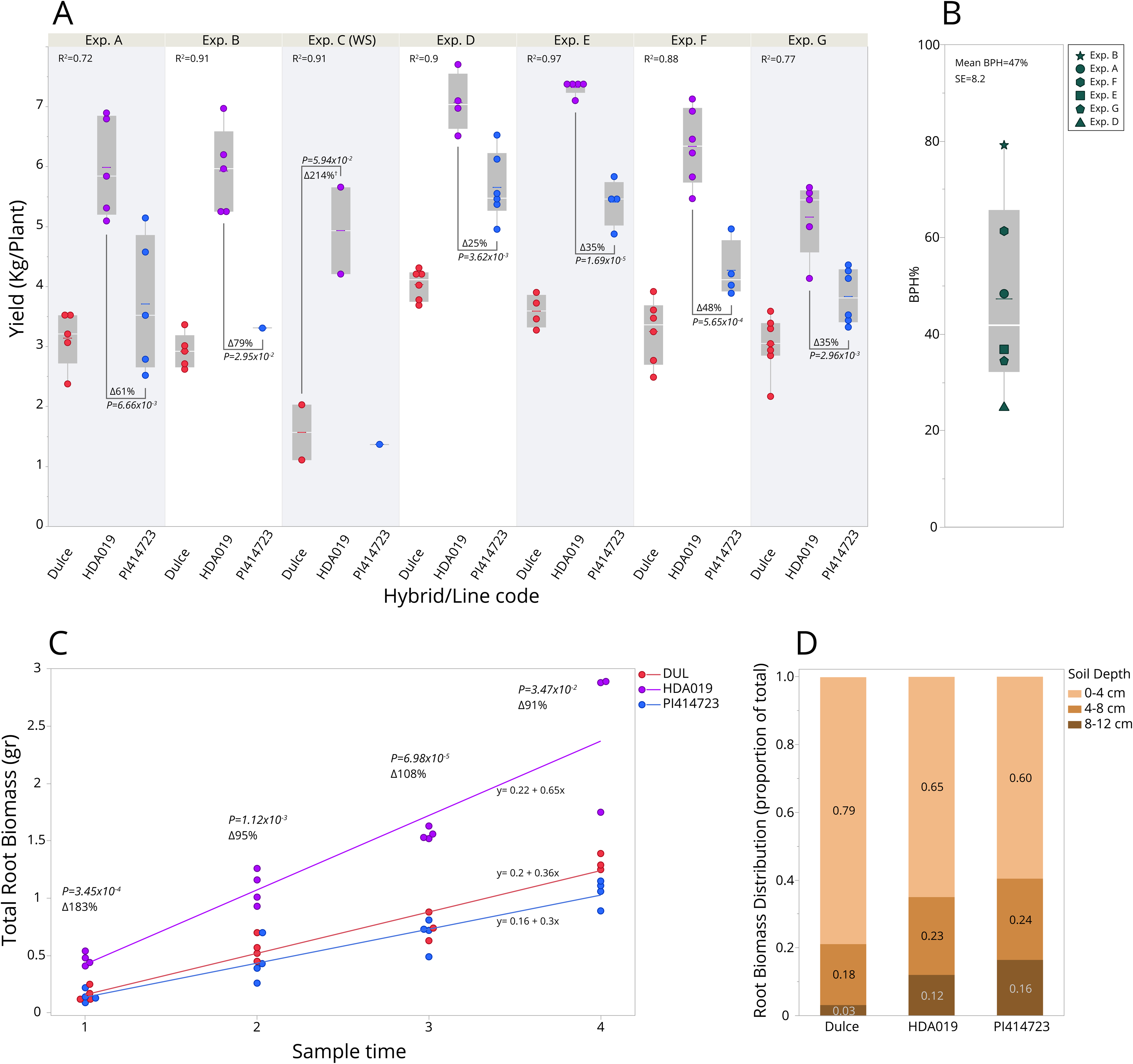
Comparative analysis of root-mediated yield and root characteristics in the HDA019 hybrid rootstock. (**A**) RMY performance of the HDA019 hybrid rootstock compared to its parental lines (PI414723 and Dulce) across six grafted field experiments conducted from 2017 to 2024. Each panel represents an individual experiment, displaying yield data (kg/plant) as boxplots with overlaid data points for HDA019 (purple), PI414723 (blue), and Dulce (red). Experiments are described in **Supplementary Table S1**. For each experiment, *R²* values, Best Parent Heterosis (BPH%), and *P*-values are provided. (**B**) Summary of Best Parent Heterosis (BPH%) across all experiments (Exp. C is excluded due to low number of replications). The boxplot shows the distribution of BPH% values with individual data points overlaid. (**C**) Comparison of root biomass between HDA019 and its parents DUL and PI414 along 4 weeks growing period. Each point represent a plant in a pot. Above each time point are the best-parent heterosis (in %) and the P value associated with this comparison. (**D**) Root biomass distribution by soil depth for HDA019 and its parental lines at the final sampling time. The stacked bar chart displays the proportion of total root biomass across three soil depth ranges: 0–4 cm (light brown), 4–8 cm (brown), and 8–12 cm (dark brown). Values within each segment represent the fraction of total root biomass at that depth.

The parental lines of HDA019 have previously been used to develop a recombinant inbred lines (RIL) population. This RIL population has been used for mapping of disease resistance and fruit quality QTLs (Harel-Beja et al. 2010; Galpaz et al. 2018). We therefore took advantage of this RIL population for genetic dissection of the RMYH of HDA019. To supplement the QTL mapping approach, and in light of the heterotic nature of yield-related traits in melon, and the significant BPH displayed by HDA019, two test-cross populations (TC-RILs) were developed by crossing each of the 78 RILs with the two parental lines, resulting in 78 DUL-TC-RILs and 78 PI414-TC-RILs (**Figure 1**). These populations expanded our ability to examine the inheritance patterns and genetic variability of the root-mediated yield variation and map QTLs under different genetic backgrounds.

### Root-mediated effects on yield, yield components and fruit quality traits in the RILs and TC-RILs populations

To characterize RMY variation, we conducted replicated yield trials using the RILs and TC-RILs populations as rootstocks with a common commercial hybrid scion (‘Glory’, a long shelf-life ‘Galia’-type hybrid variety), over two consecutive years (RILs and DUL-TC-RILs were tested in both years and PI414-TC-RILs was tested only in the second year). The two parental lines and the F_1_, as well as ‘Glory’ grafted on itself were used as genetic controls in these experiments.

In both years and across the three populations, RMY displayed normal distribution, confirming the expected quantitative nature of this trait (**Figure 3A**). In accordance with the results from the HDA experiments (Dafna et al. 2021), ‘Glory’ grafted with DUL rootstock was at the lowest end of the yield distribution, ‘Glory’ grafted with PI414 rootstock was in the middle range, and ‘Glory’ grafted with the F_1_ hybrid (HDA019) was at the highest end of the RMY range (**Figure 3A**).

**Figure 3:**
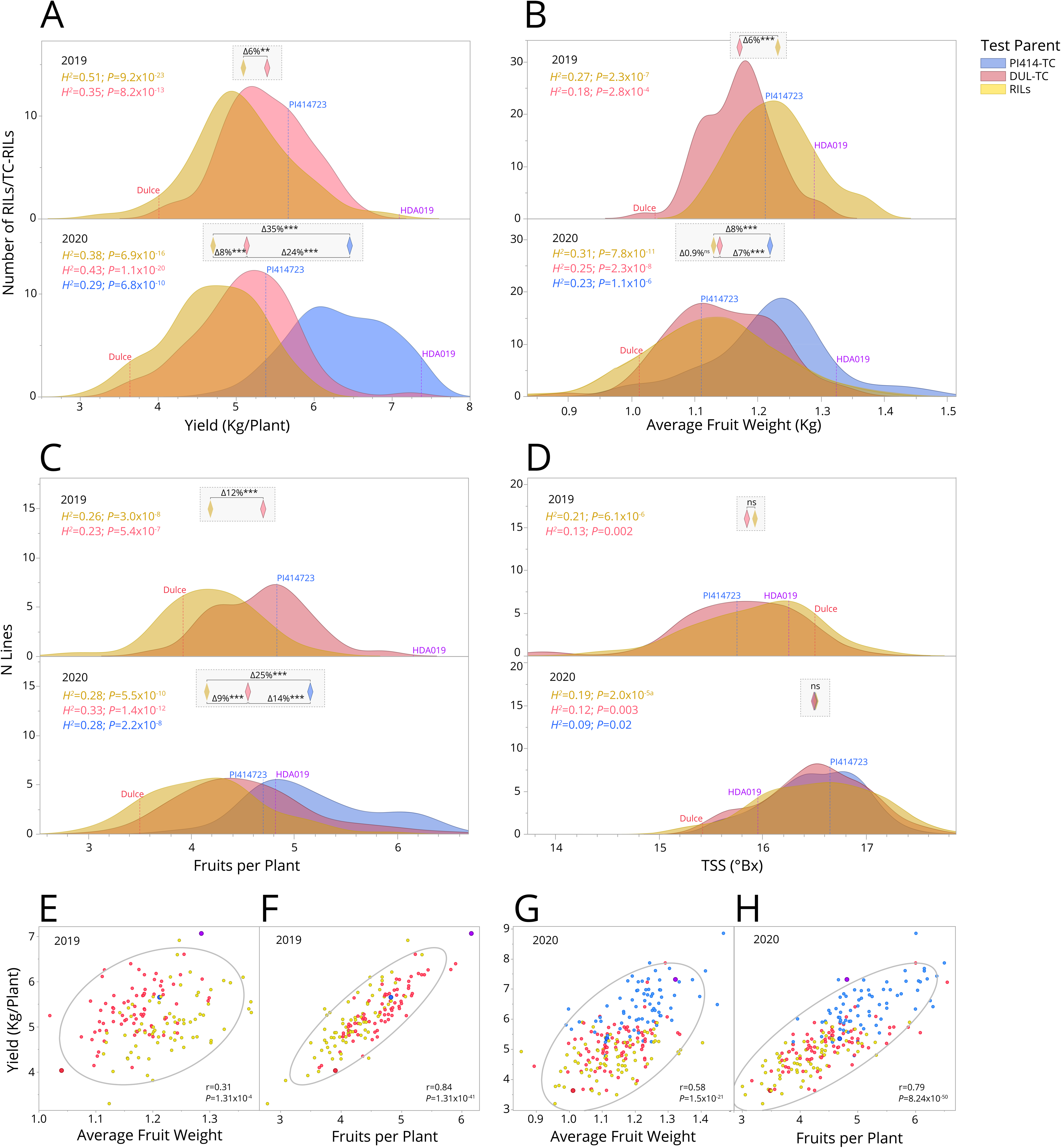
Trait distributions and correlations in grafted PI414 × DUL RIL and TC-RIL populations. (**A–D**) Distribution of RMY and fruit traits in grafted PI414 × DUL Recombinant inbred lines (RILs) and test-cross RILs (TC-RILs) over two growing seasons (2019 and 2020). Traits analyzed include (**A**) Yield (kg/plant), (**B**) Average fruit weight (kg), (**C**) Fruits per plant, and (**D**) Total soluble solids (TSS, °Brix). Each panel displays density plots for the RIL population (yellow) and both TC-RIL populations (PI414-TC in blue, DUL-TC in pink). Parental lines (Dulce and PI414723) and the F_1_ hybrid (HDA019) are also indicated. Rectangular insets provide Δ values representing statistical comparisons between sub-population means, with asterisks denoting significance levels. (**E–H**) Correlation analyses between yield and its components for the 2019 and 2020 growing seasons. Scatter plots display relationships between yield and average fruit weight (**E, G**) and yield and fruits per plant (**F, H**), with color-coded points representing RILs and TC-RILs. Ellipses indicate confidence intervals, and *r*-values (Pearson’s correlation coefficients) and *P*-values are provided.

A significant genetic background effect was found; ‘Glory’ grafted with the PI414-TC-RILs rootstocks yielded on average 35% more than the parallel grafting with the inbreds (RILs) rootstocks (*P*=6.0 × 10^-32^); ‘Glory’ grafted with the DUL-TC-RILs rootstocks yielded on average 8% more than the parallel grafting with the inbreds (RILs) rootstocks (*P*=1.4 × 10^-4^, **Figure 3A**).

RMY range (on entry-mean basis) within each sub-population was ∼2-fold: 3.2-5.9 Kg/Plant in the inbred rootstocks, 3.6-7.9 Kg/Plant in the DUL-TC-RILs rootstocks and 4.8-8.9 Kg/Plant in the PI414-TC-RILs rootstocks (data for 2020; **Figure 3A**). The proportion of yield variation explained by root-mediated genetic effects was significant in all these experiments and genetic backgrounds (**Figure 3A**). For example, broad-sense heritability in the RILs population was *H^2^*=0.51 *(P*=9.2 × 10^-23^) in 2019 and *H*^2^=0.38 *(P*=6.9 × 10^-16^) in 2020. These results further assured the suitability of this bi-parental population for genetic mapping of RMY.

To understand better the basis for the observed RMY variation, we collected phenotypic data across all experiments also on its components; number of fruits per plant (FN) and average fruit weight (AFW). AFW values displayed similar distributions across the 3 populations (**Figure 3B**). In the RILs, the range was between 1.11-1.37 and 0.85-1.40 kg/fruit in 2019 and 2020, respectively. Significant genetic effects (*H^2^*) were calculated in both years in all the populations, with values ranging between 0.18 and 0.31 (**Figure 3B**). Number of fruits per plant (FN) also displayed similar distributions across the 3 populations (**Figure 3C**). In the RILs, the range was between 2.78-5.34 and 3.00-5.89 fruits/plant in 2019 and 2020, respectively. Significant genetic effects (*H^2^*) were calculated in both years in all the populations, with values ranging between 0.23 and 0.33 (**Figure 3C**). As in the HDA experiments (Dafna et al. 2021), RMY variation across the RILs and TC-RILs was explained by variation in its components, root-mediated FN and root-mediated AFW. Strong positive correlation was found between root-mediated FN and RMY in a combined analysis of the populations (*r*=0.84 in 2019 and *r*=0.79 in 2020; **Figure 3F, H**) and a weaker positive correlation is shown with root-mediated AFW (*r*=0.31 in 2019 and *r*=0.58 in 2020; **Figure 3E, G**). Collectively, the meaning of these analyses is that higher RMY is explained by increase in the number of fruits as well as in their average weight.

We also collected data on total soluble solids (TSS) as a strong indicator of fruit sweetness, and a major quality determinant in melon breeding. TSS values did not differ between the 3 populations (**Figure 3D**), ranging in the RILs between 14.4-17.1 and 15.3-17.5 Bx° in 2019 and 2020, respectively. Marginal genetic effects (*H^2^*) were calculated in both years in all the populations, with values ranging between 0.09 and 0.21 (**Figure 3D**), indicating that while rootstocks genotype affect yield and its components, it does not affect fruit sweetness.

### Mapping and prioritization of root-mediated yield QTLs

To genetically dissect the yield variation and map QTLs that contribute to the observed RMY improvement of HDA019 F_1_ hybrid, we performed QTL mapping analysis. The PI414 × DUL RILs population was previously genotyped with 44K informative SNPs (Galpaz et al. 2018) which were binned into 1,763 bin-markers, evenly distributed throughout the melon genome (Oren et al. 2020). Linkage analysis was performed separately for the RILs and TC-RILs in each experiment.

Overall, and in line with the quantitative nature of RMY, effects detected through the QTL mapping analyses were moderate. We initially identified 20 genomic regions that showed multiple marker-trait associations across different years and sub-populations, which independently displayed a non-adjusted significance of *P*<0.05 (for each such region, peak QTL marker is presented in **Supplementary Table S3**). To obtain broader perspective and prioritize the observed association signals affecting RMY, we undertook a unified analysis approach by analyzing the set of 20 associated QTL regions across 4 experiments—each combination of background and year was treated as a unique experiment, PI414-TC-RILs 2020 dataset was excluded from this analysis due to the significant shift toward higher RMY of this background (**Figure 3A**)—aggregating to 80 trait-QTL tests (**Supplementary Table S3**). The criteria for selecting consistent RMY QTLs were established based on an RMY allelic effect surpassing 5% in the same direction in at least three experiments, resulting in the identification of 9 QTL regions showing this robust pattern, as illustrated in the QTL Venn diagram, showing the intersections of such common significant QTLs across experiments (**Figure 4A, and Supplementary Table S3**). For each of the 9 QTLs we then analyzed the combined genotype effect across the 5 experiments (2-3 populations and two years), and after this phase we selected the five most consistent QTLs (hereafter, ‘*Top5*’), located on chromosomes 1, 2, 6, 10 and 11 for further investigation (**Figure 4B**). The power of the combined analysis for selecting reliable and consistent RMY QTLs is demonstrated in **Figure 4C**, displaying allelic means of each QTL across 5 populations/experiments. The consistency of the selected QTLs with moderate RMY effect of only ∼6% is shown and translated to significant genetic effect statistics in a unified two-way QTL × population-experiment analyses (**Figure 4D**, *P (G)* values of 10^-5^ – 10^-7^).

**Figure 4:**
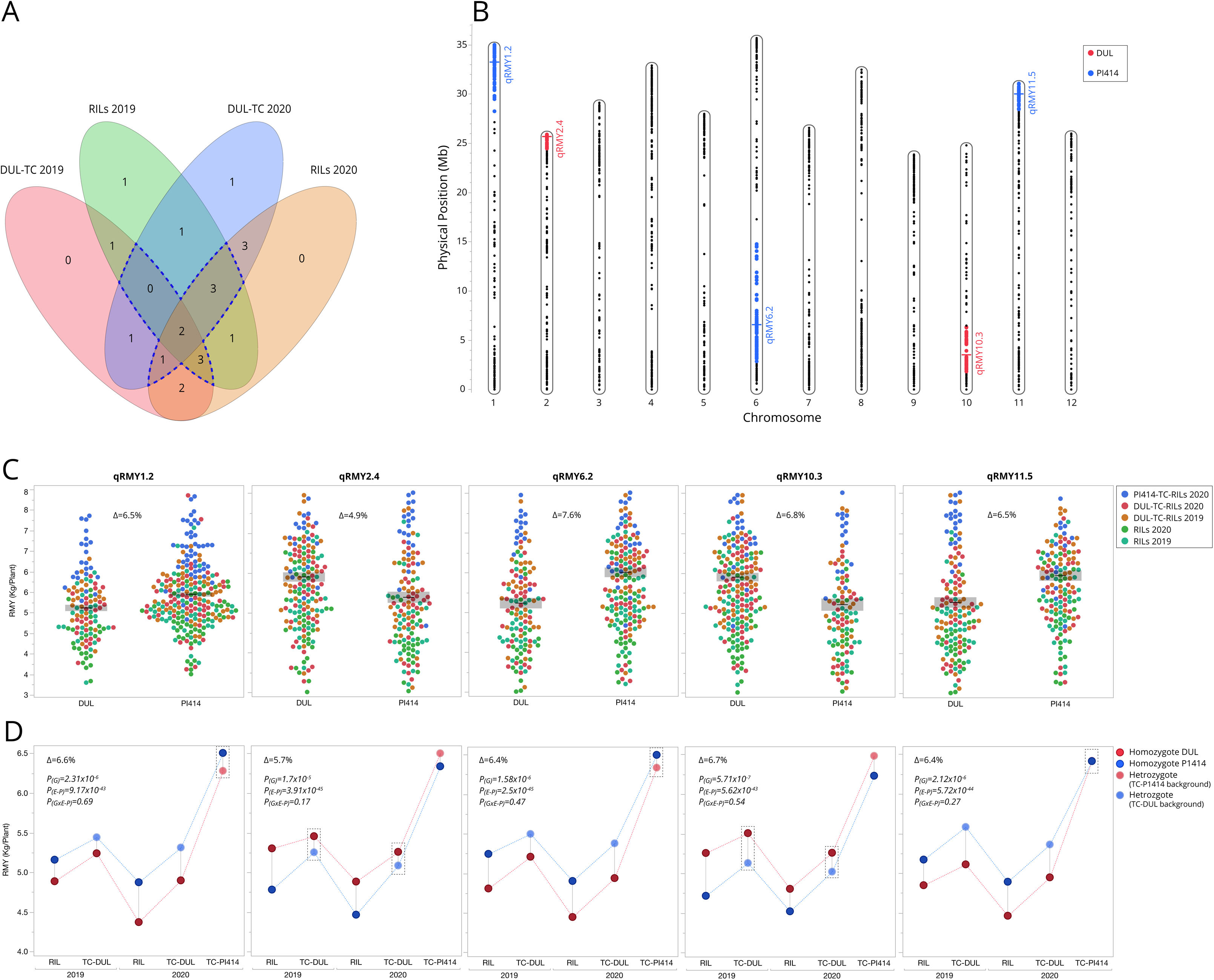
Mapping and prioritization of root-mediated yield (RMY) QTLs. (**A**) Venn diagram illustrating the overlap of nine RMY QTLs across different sub-populations and years. Each experiment (RILs 2019, RILs 2020, DUL-TC 2019, and DUL-TC 2020) was treated as a separate dataset, and QTLs were retained if their yield effect exceeded 5% in at least three experiments (dashed line). The numbers in each region indicate the count of shared QTLs among datasets. (**B**) Genomic positions of the five selected QTLs based on physical position (Mb) across 12 melon chromosomes. The selected QTLs (qRMY1.2, qRMY2.4, qRMY6.2, qRMY10.3, and qRMY11.5) are highlighted in blue (where PI414 allele is favorable) and red (where DUL allele is favorable) (**C**) Combined RMY effect of the ‘Top5’ QTLs across different populations and years. (**D**) Allelic RMY effects of the ‘Top5’ selected QTLs in each of the experiments. The Δ values and *P*-values represent a combined two-way ANOVA (Genotype × Experiment-population). Comparisons are shown for RILs, DUL-TC, and PI414-TC populations. *P*(G) - *P* value for QTL genotype effect. *P*(E-P) - *P* value for Experiment-population effect. *P*(G × E-P) - *P* value for the interaction between QTL genotype × Experiment-population. Dashed rectangles highlight comparisons between heterozygotes and homozygotes to the favorable alleles.

### Allelic effect and mode-of-inheritance of ‘TOP5’ RMY QTLs

Across the ‘*Top5*’, in three QTLs the favorable allele is from PI414 (qRMY1.2, 6.2 and 11.5, **Figure 4C-D**), and the RMY increasing allele in the two other QTLs (qRMY2.4 and 10.3) is from DUL. While there is significant difference in the performance of the parental lines *per se* and their test-cross populations (**Figures 2A**, **3A**), the fact that favorable RMY QTL alleles are contributed by both parents is in line with the transgressive segregation in the RILs and TC-RILs populations and the extensive RMY heterosis of *HDA019*.

As described above, the effect of each of the ‘*Top5*’ RMY QTLs was evaluated and found to be consistent across 5 experiments representing 3 different populations and genetic compositions. While in none of the populations there is representation of the 3 genotypic categories of a QTL (homozygote DUL, heterozygote and homozygote PI414), the two testcross populations provide insight into the comparison between heterozygotes and homozygotes DUL (in the DUL-TC-RILs), or homozygotes PI414 (in the PI414-TC-RILs), and as such complement the comparisons between DUL and PI414 homozygotes, at the RILs population. One of the obvious questions of this study was whether the heterotic effect of HDA019 is explained also by overdominant QTLs in the derived segregating populations. The fact that in none of the ‘*Top5*’ QTLs, heterozygotes displayed higher RMY compared to the homozygotes for the favorable allele (**Figure 4D**; dashed rectangles) indicates that their mode-of-inheritance (MOI) is not over-dominance. qRMY1.2, 6.2 and 11.5 display dominant effect as the effects of heterozygotes for the favorable (PI414) allele in the DUL-TC-RILs is similar to the effect of these QTLs in the homozygote RILs (**Figure 4D**). qRMY10.3 also displays dominant effect as the difference between homozygotes for the favorable (DUL) allele and heterozygotes in the DUL-TC-RILs is similar to the effect of this QTL in the homozygote RILs (**Figure 4D**). qRMY2.4 displays additive MOI trend as the effect in the DUL-TC- RILs is smaller compared to the effect in the RILs, in both years (**Figure 4D**).

### Interactions and combined effects of RMY QTLs

In line with our interest in the ‘real-life’ breeding implication of this study, the subsequent phase focused on evaluating the interactions between RMY QTLs, and more specifically the combined effect of pairs or triplets of QTLs, with an emphasis on the potential for pyramiding these QTLs. Examples for the combined effect of three QTL-pairs from the ‘*Top5*’ are provided in **Figure 5A-C**, where the non-significant epistasis results in a cumulative pattern of QTL additivity. To explore the RMY QTL epistasis in a more comprehensive way and provide a broader view on the mostly additive (non-epistatic) pattern between QTLs, we expanded the analysis to a broader (less stringent) set of 9 RMY QTLs that were defined from the initial 20 QTL-trait associations (**Supplementary Table S3**). For this set, we first calculated the expected—non-epistatic—effects of 36 QTL-pairs and 84 QTL-triplets as the sum of their singular effects. The expected effects were then plotted against the actual observed performance of these QTL-pairs and triplets in the RILs and DUL-TC-RILs over two years (**Figure 5D-G; Supplementary Table S4**). A significant linear relation was found between observed and expected (*R^2^* = 0.78 and 0.70 in the RILs in 2019 and 2020, respectively, **Figure 5D-E**) indicating the robustness of the additive component in QTL interactions. Still, most of the points are located below the observed = expected diagonal, suggesting that less-than-additive is a relevant pattern, in particular for combinations of large-effect QTLs. General less-than-additive epistasis is also confirmed by the slopes of regression lines for the expected versus observed of the QTL-pairs, which are significantly lower than 1.00 (0.77 and 0.67 in the RILs in 2019 and 2020, respectively, **Figure 5D-E**). To assess the potential RMY enhancements of multi-QTL combinations, we tested differences between RILs genotypes carrying favorable alleles in 1-5 of the ‘Top5’ QTLs. In both years, we show RMY increase associated with higher number of QTLs, and RILs carrying favorable alleles in 4 and 5 QTLs had significantly higher RMY compared to RILs with lower number of QTLs (**Figure 5H-I**).

**Figure 5:**
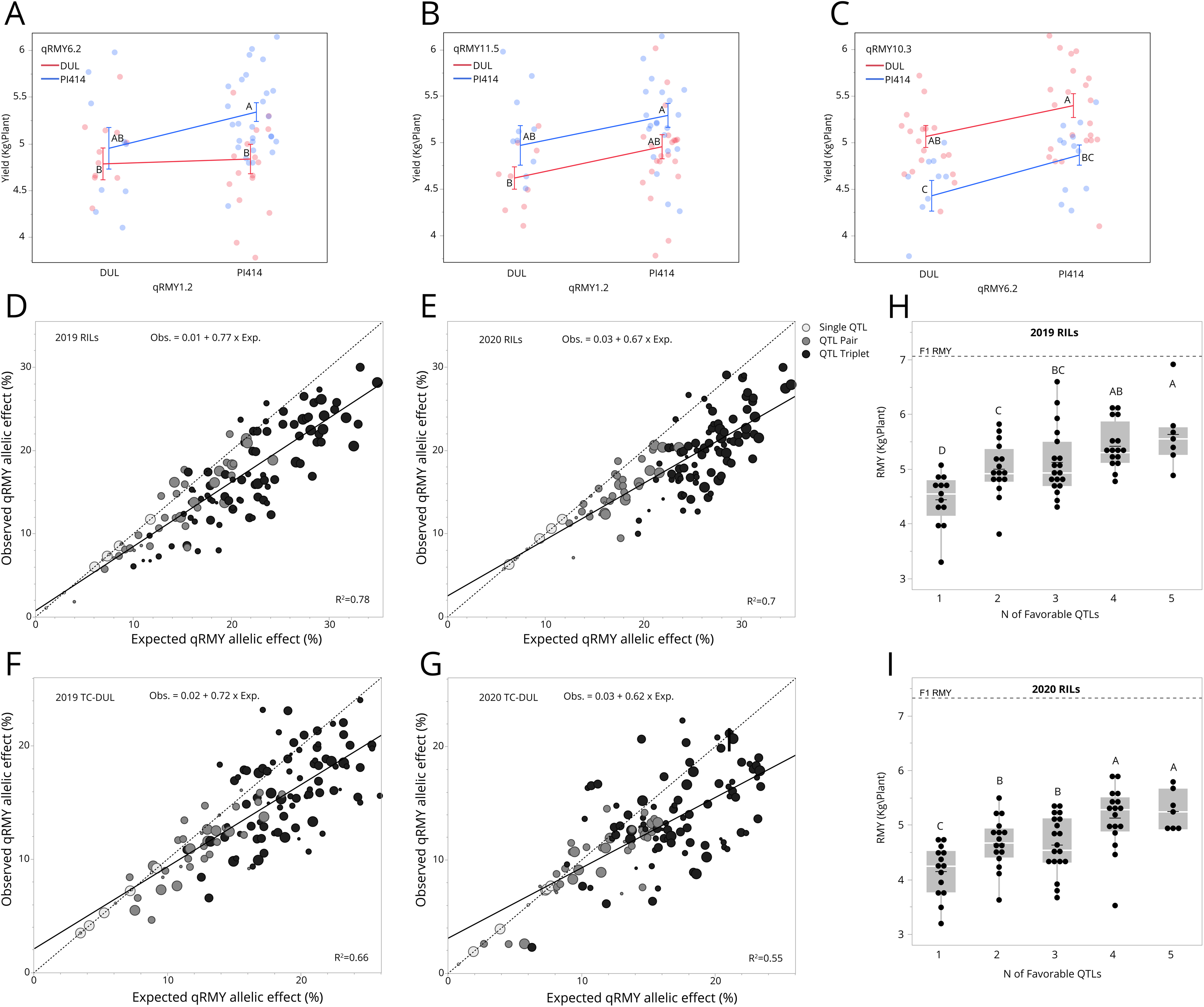
Interactions and combining patterns of root-mediated yield (RMY) QTLs. (**A–C**) Examples for analyses of QTL pair interactions in the 2019 RILs experiment. Values not connected by a common letter are significantly different at P<0.05. (**D-G**) Correlation between expected and observed QTL effects for single QTLs, QTL pairs, and QTL triplets for 9 QTLs across the RILs and DUL-TC-RILs populations. Dashed diagonal indicates expected = observed, and the linear regression function is displayed in each panel. Circle size denotes the proportion of ‘Top5’ selected QTLs within each combination. (**H-I**) five-QTL stacking effect in the RIL population in 2019 (**H**) and 2020 (**I**). A dashed line at the top of each panel marks the F_1_ hybrid’s performance as a reference.

### Validation of QTL contribution to root-mediated yield using QTL-backcross lines

Analysis of RMY in the RILs and TC-RILs demonstrated that the yield variation is mediated by small effect QTLs, which are inherently more challenging for characterization. To empirically validate the specific contribution of QTLs for RMY in a field setting, QTL-backcross (QTL-BC) lines were advanced through marker-assisted backcrossing of the 5 selected QTLs into their respective non-favorable parental backgrounds (**Figure 1** and Methods section). The QTL-BC lines, harboring favorable RMY allele in the QTL region, were evaluated in a segregating population setup over 2-3 generations (BC_1_F_2_, BC_3_F_2_ and BC_3_F_3_). In each BC generation, comparison was performed between plants homozygotes for the favorable and non-favorable QTL-alleles, with the recurrent parents used as another genetic reference. In the BC_3_ generations of both PI414 and DUL backgrounds, RMY of the non-favorable segregants was not significantly different from that of the corresponding recurrent parent (**Figure 6A-B**), indicating on the efficacy of the marker assisted BC, as also visually observed by similarity in fruit characteristics (**Supplementary Figure S2**). In three of the QTLs (qRMY2.4, qRMY6.2, qRMY11.5) we showed significant 10%-16% RMY increase associated with the favorable allele in the advanced BC_3_ generation (**Figure 6D-F**). Interestingly, in QTL qRMY1.2 that showed a consistent Δ6.6% effect in the RILs and TC-RILs (**Figure 4C-D**), and a Δ37% RMY increase in the BC_1_F_2_ generation (**Figure 6C**), the effect was abolished in the BC_3_F_2_ and BC_3_F_3_ generations (**Figure 6C**), suggesting a possible positive epistasis that was removed through the backcrossing process, as an explanation for the loss of this QTL effect. qRMY10.3, which showed a consistent RMY effect in the RILs and TC-RILs (**Figure 4C-D**), was tested only at the BC_3_F_3_ generation and displayed a non-significant 5.6% RMY increase by the favorable DUL allele (**Figure 6G**). Further analysis of this QTL will be required. Collectively, these analyses provide important independent validations for the effects of these QTLs on root-mediated melon yield and are, therefore, an important step towards focused fine-mapping and broader investigation of their independent and combined breeding values.

**Figure 6:**
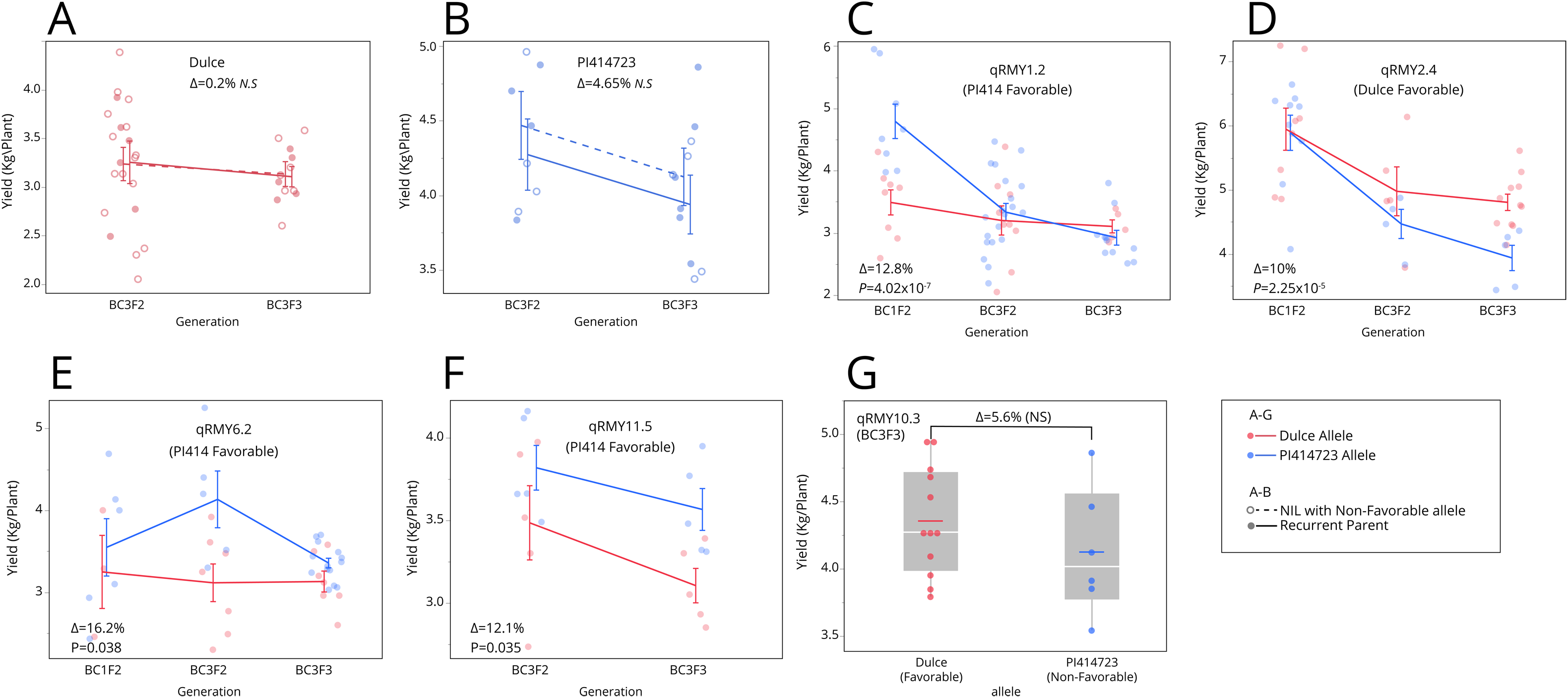
Validation of ‘Top5’ QTL RMY Effects in QTL-BC lines in grafted field experiments. (**A**) Yield performance of NILs carrying the non-favorable allele at qRMY1.2, qRMY6.2 and qRMY11.5, relative to their recurrent parent background, Dulce (**B**) Yield performance of NILs carrying the non-favorable allele at qRMY2.4 and qRMY10.3, relative to their recurrent parent background, PI414723. (**C-F**) Yield performance of QTL-BC lines homozygotes for the favorable or non-favorable allele for qRMY1.2, qRMY2.4, qRMY6.2, and qRMY11.5 across successive backcross generations. The Δ values and *P*-values represent a combined two-way analysis (yield × experiment). (**G**) Yield comparison of NILs carrying the favorable allele versus those with the non-favorable allele for qRMY10.3.

### Potential of HDA019 in melon commercial production

In a previous study, we demonstrated that hybrid rootstocks significantly outperform rootstocks of inbred lines in mediating fruit yield (Dafna et al. 2021). In the current study we genetically dissected RMY of one of the best performing hybrids, HDA019. In addition to extensive validation of the RMYH of this hybrid compared to its parents (**Figure 2A-B**), from a breeding standpoint it is important to evaluate the performance of this hybrid rootstock under multiple scion varieties and compared with relevant commercial rootstocks used in melon production. **Figure 7** is summarizing the performance of HDA019 rootstock across 22 independent comparisons that represent different experiments (reflecting different growing conditions, **Supplementary Table S1**) and 5 different scion genotypes. In each of these experiments, RMY performance of the HDA019 rootstock was compared to the respective non-grafted scion genotype and to a commercial Cucurbita hybrid rootstock (TZ). In some of the experiments, commercial melon rootstocks (MRS2 and Aroma) were also used as references. The results are presented as percentage difference from each of the non-grafted scion varieties (detailed in **Supplementary Table S5)**. While there is variation in the relative performance of HDA019 across the different experiments and scions, a consistent advantage of this hybrid rootstock over both the non-grafted (mean Δ24%) and the commercial Cucurbita rootstock is shown. In Experiments K, N, O and P, under the ‘Glory’ scion, HDA019 also outperformed MRS2 and Aroma, two commercial melon rootstocks. These results reflect the practical potential of HDA019 and additional melon hybrid rootstocks that we previously tested (Dafna et al. 2021) for enhancing melon production under variable field conditions.

**Figure 7:**
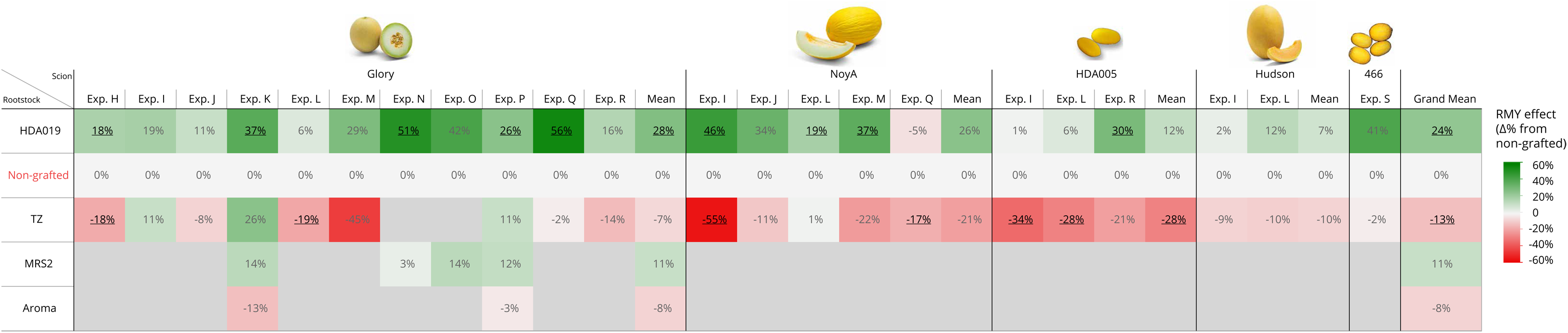
Root-mediated yield performance of HDA019 rootstock compared to non-grafted and commercial Cucurbita and melon rootstocks. Values represent the percentage difference in RMY relative to the non-grafted control for each scion variety. Glory is a yellow-rind Galia-type variety, NoyA (Noy-Amid) is a yellow-canary-type accession, HDA005 is a small fruited experimental hybrid, Hudson is an Ananas type variety, 466 is an experimental small-fruited inodorous variety.

## Discussion

### Dissection of a yield-enhancing melon hybrid rootstock

This study presents a genetic dissection of root-mediated yield heterosis (RMYH) in melon, expanding upon our previous findings (Dafna et al. 2021). By grafting a segregating recombinant inbred line (RIL) population as rootstocks, the study characterizes the polygenic nature of RMYH and its inheritance patterns. The experimental system was based on a RIL population derived from HDA019, a hybrid rootstock that consistently exhibited best-parent heterosis (BPH) for yield across multiple environments. The RIL population previously developed from this F_1_ hybrid (Danin-Poleg et al. 2002; Harel-Beja et al. 2010) enabled a systematic investigation of the inheritance patterns and genetic architecture of RMY variation. Prior experiments identified several melon hybrids—including HDA010, HDA016, HDA019, and HDA100—as high-performing hybrid rootstocks, with HDA019 selected for further genetic dissection due to its strong and stable heterotic effects (RMY BPH of 25–79% across years and growing conditions; **Figure 2A, B**; (Dafna et al. 2021).

In addition to its superior yield-related field performance, the heterotic rootstock HDA019 also exhibited a distinct root biomass accumulation pattern compared to its parental lines. Preliminary root system analyses indicated that this hybrid rootstock had greater total root biomass and a more balanced root distribution across soil depths (**Figure 2C, D**), potentially optimizing resource acquisition. While these findings are based on a limited sample size and require validation in broader additional experiments, they align with previous studies showing that deeper and more evenly distributed root systems enhance water and nutrient uptake efficiency, ultimately improving plant productivity (Lynch 2013, 2019; Koevoets et al. 2016).

### Genetic architecture of root-mediated yield inferred from analysis of melon RILs and TC-RILs populations

In both years and across all three populations, root-mediated yield (RMY) exhibited a normal distribution, consistent with its quantitative nature (**Figure 3A**). The RILs and TC-RILs populations captured a significant heritable RMY variation, but interestingly, no individual line in the population reached the performance of the F_1_ hybrid rootstock (**Figure 3A**). This observation, along with the finding that favorable alleles for the main RMY QTLs were contributed by both parents (**Figure 4**), supports the hypothesis that heterosis for root-mediated yield in HDA019 arises from the cumulative effect of multiple genes, rather than from contribution of major quantitative trait loci. Similar findings have been reported in other crops, where yield-related traits are largely controlled by multiple small-effect loci dispersed throughout the genome (Buckler et al. 2009; Schnable and Springer 2013; Huang et al. 2015).

The test-cross populations (TC-RILs) were designed to expand the genetic analysis to better define the mode of inheritance of RMY QTLs and examine whether we can detect QTLs that display locus-specific heterosis that contributes to RMY, as described in other crop species (Hua et al. 2003; Semel et al. 2006; Krieger et al. 2010; Guo et al. 2014). Each RIL represents a unique mosaic of genome-wide recombination pattern in a homozygous state, while its respective TC-RIL captures also the heterozygous state at those same loci, enabling the detection of overdominant effects at specific genomic regions. Analysis of mode of inheritance of the ‘Top5’ RMY QTLs in the TC-RILs showed that in none of them, the QTL at its heterozygote state outperformed the corresponding homozygotes for the favorable allele (**Figure 4D**). These results support a dominant complementation or QTL epistasis models (Xiao et al. 1995; Lippman and Zamir 2007; Garcia et al. 2008) as the more probable and prevalent modes to explain the RMY heterosis of HDA019.

### Root-mediated yield variation is mainly explained by the number of fruits per plant

Further examination of yield components revealed that higher RMY was primarily driven by an increase in fruit number rather than the average fruit weight (**Figure 3B, C, E-H**). Correlation analyses confirmed that both traits were positively associated with yield, though fruit number per plant (FN) exhibited a stronger correlation (*r* = 0.84–0.79 in 2019 and 2020, respectively; **Figure 3F, H**) compared to average fruit weight (AFW) (*r* = 0.31–0.58 in 2019 and 2020, respectively; **Figure 3E, G**). This suggests that rootstocks influence yield primarily by affecting regulation of fruit setting rather than fruit size. These results are consistent with the correlations calculated between yield and its component in our previous study on RMY heterosis in a diallele population (Dafna et al. 2021).

This pattern aligns with the hypothesis that yield increase in hybrid rootstocks results from improved overall plant fitness and adaptability to growing conditions. While fruit size is highly heritable and genetically constrained by the scion genotype, fruit number is more responsive to external and internal inputs, including resource availability, allowing plants with enhanced root function and resource acquisition to sustain a greater fruit load (Marcelis et al. 2004; Barzegar et al. 2018).

### The potential for melon yield enhancement through grafting with hybrid rootstocks

The results of the current study and our previous broad investigation of inheritance of root-mediated yield in melon (Dafna et al. 2021), both point out that heterosis is a prominent factor contributing to the yield increase by rootstocks. The indications that RMY heterosis is explained by multiple small-effect loci acting in a cumulative dominant or additive manner is reminiscent for the contribution of yield heterosis demonstrated in major crops (Paril et al. 2024). Hybrid-breeding strategies in such crops are primarily based on the definition of distinct heterotic groups and improvement of inbred lines through pedigree selections within the defined groups and based on early-generation test-cross performance. Experimental hybrid combinations are then created as test-crosses between lines from the different heterotic groups. In melon (and other vegetables), the breeding strategy is different and mostly based on breeding within narrow market-type-defined groups to maintain strict fruit characteristics. This practice is limiting the potential utilization of heterosis for yield improvement. Our current and past results (Dafna et al. 2021) on the importance of RMY heterosis, support the proposal of separate breeding workflows for rootstock and scions in melon, which will implement different methodologies and allow each product to be developed using the optimal breeding strategy (**Figure 8**). The physical and conceptual separation of breeding above (scion) and underground (rootstock) varieties ensures maximal flexibility to breed heterotic melon rootstocks by leveraging heterotic-group-oriented breeding strategy (**Figure 8B**), and on the same time maintain the classical melon breeding methodology to develop scion varieties with strict, market-type-defined, high fruit qualities (**Figure 8A**). The heterosis-oriented rootstock breeding promote utilization of broad germplasm diversity, including exotic melons and cultivar-groups that are not routinely used in conventional melon breeding (**Figure 8B**). This approach allows efficient and simple introgression of soil pathogen resistances into the rootstock varieties, and therefore simplifying the scion breeding by eliminating the need to breed for these traits. The optimal combination between scion and rootstock will be the final phase of selecting grafted varieties adapted to specific growing conditions and market demands (**Figure 8C**). As demonstrated in the current study, genetic mapping and genotypic selection of RMY QTLs is feasible for stacking favorable alleles to improve rootstock inbred lines (**Figure 5**) as a complementary route to the utilization of heterosis for RMY improvement. The ‘Top5’ RMY QTLs that we investigated displayed significant cumulative pattern when combined (**Figure 5 A-C, H, I**). However, on a broader level, a less-than-additive mode of interaction between QTLs was found to be significant (**Figure 5 D-G**). Such a diminishing effect of QTL epistasis was also previously described for TSS and yield-related traits in tomato (Eshed and Zamir 1996; Gur and Zamir 2015) and melon (Tzuri et al. 2025).

**Figure 8:**
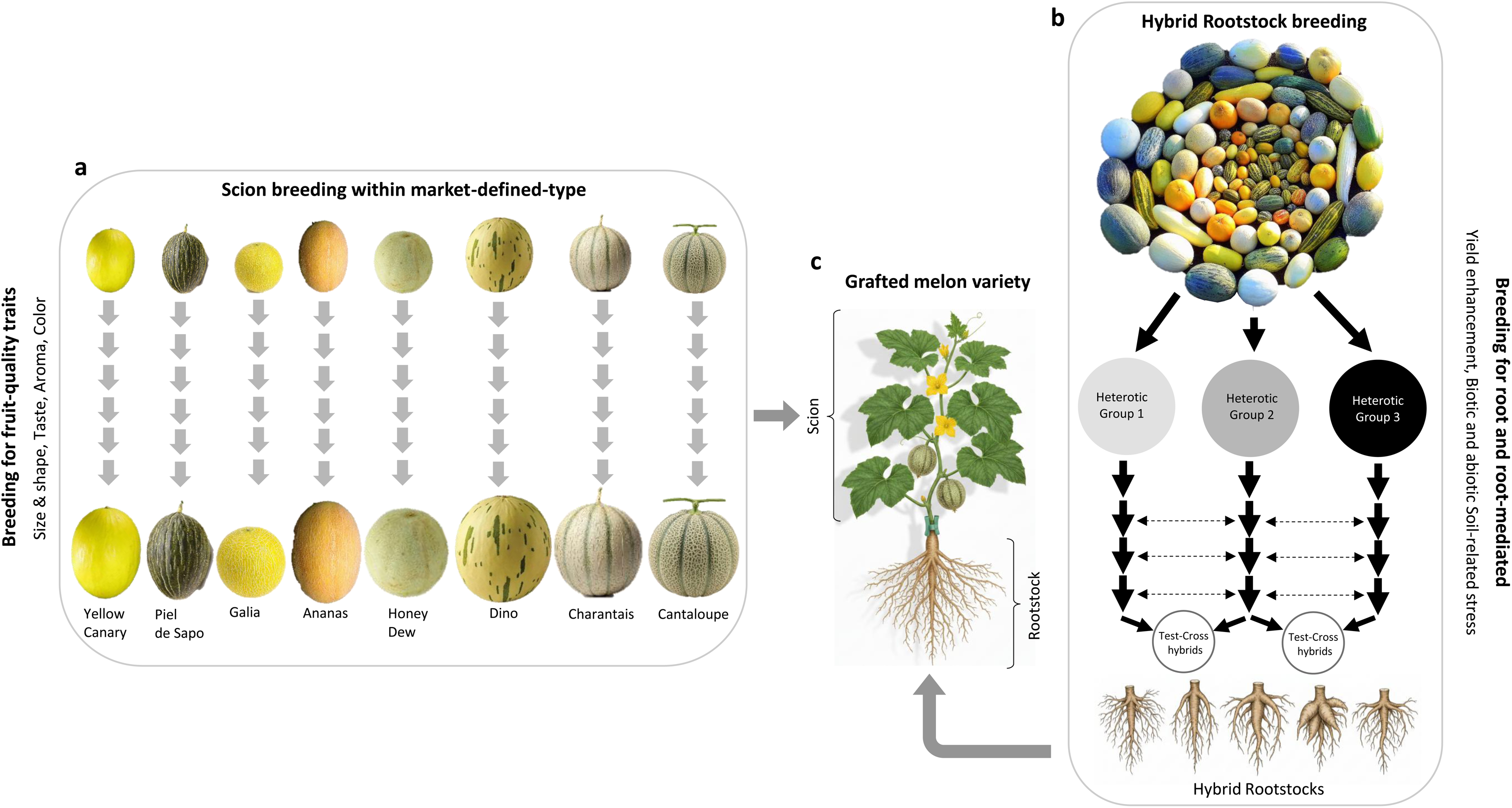
Schematic illustration of the split-breeding approach, demonstrating: (**A**) Scion breeding – focused on plant health and fruit quality traits. (**B**) Hybrid rootstock breeding– focused mainly on heterosis driven root-mediated yield enhancement. (**C**) Testing under different soil types and growing conditions and selecting best-performing grafted scion-rootstock combinations.

### The challenges and potential of improving crop yield using root traits

Root biology has increasingly attracted scientific attention in recent years, being recognized as a promising avenue for enhancing plant productivity under both favorable and stress-induced conditions (Bagale et al. 2025). Nonetheless, the majority of genetic studies in model and crop species have largely adopted a foundational perspective, concentrating primarily on the examination of root development and variation in root system architecture (Paez-Garcia et al. 2015; Tracy et al. 2020), rather than undertaking of challenging direct investigations into functional variation within root systems. In this regard, grafting represents a powerful experimental and practical tool, enabling the separation—and to a large extent, independent investigations—of shoot (scion) and root (rootstock) functional properties and their effects on whole-plant performance under field conditions. While there is expanding use of grafted plants in Cucurbitaceous and Solanaceous vegetable crops (Bahadur et al. 2024), including reports on beneficial rootstocks (Edelstein et al. 2004; Gong et al. 2022; Ingram et al. 2022; Davis et al. 2024; Ndlovu et al. 2025), there is still very limited research into the genetics underlying yield enhancing rootstocks. In tomato, only few studies have taken a QTL mapping approach to genetically dissect the variation in root-mediated scion traits using grafting (Estañ et al. 2009; Gur et al. 2011; Asins et al. 2020; Kabas et al. 2025). To the best of our knowledge, the current study is the first genetic mapping analysis of rootstock-mediated yield variation in a cucurbit crop. The complexity of genetically dissecting the contribution of roots to yield variation has, thus far, only exposed a part of the genetic architecture, in the form of low resolution QTLs. Discovery of causative genes underlying root-mediated yield QTLs remains a future challenge, and the advanced QTL-BC lines developed in the current study are potential substrates towards such goal. A systematic, focused, genetic research of rootstocks in grafting amenable crops could be a rewarding effort, as it has the potential to uncover yield- and stress-tolerance-related genes and genetic variants affecting the hidden, underground, target of plant breeders.

## Supplementary data

The following supplementary data are available online.

**Supplementary Table S1:** List of Experiments

**Supplementary Table S2:** List of primers used for genotypic selection of ‘TOP5’ RMY QTLs

**Supplementary Table S3:** List of RMY associations detected in the RILs and TC-RILs populations

**Supplementary Table S4:** Observed vs. expected effects of RMY QTL combinations

**Supplementary Table S5:** HDA019 performance vs. commercial rootstocks

**Supplementary Fig. S1:** Distribution of root biomass by soil depth across 4 time-points in PI414, DUL and their F_1_ hybrid (HDA019).

**Supplementary Fig. S2:** Fruit images of recurrent parent (PI414) and two QTL-BC lines at the BC_3_F_3_ generation

## Statements and Declarations

## Acknowledgments

We thank Fabian Baumkoler, Zohar Ben-Simhon and the farm team at Newe Ya’ar for technical assistance in setting the field trials and for plant maintenance.

## Author contributions

AG and AD: conceptualization; AD, GT, EO, TI, IH, GP and AG: investigation; AD, EO and AG: formal analysis; AD and AG: writing - original draft; AD and AG: visualization; All authors read and approved the final manuscript.

## Data availability statement

The data supporting the findings of this study are available within the paper and within its supplementary materials published online.

## References

Alarcón AL, Gómez-Bellot, María José, Bernabe, Antonio José, et al (2020) Changes in root architecture and productivity of melon (Cucumis melo L. cv. Hispano Nunhems) promoted by Glomus iranicum var. tenuihypharum. J Hortic Sci Biotechnol 95:364–373. 10.1080/14620316.2019.1681906

Asins MJ, Raga MV, Torrent D, et al (2020) QTL and candidate gene analyses of rootstock-mediated tomato fruit yield and quality traits under low iron stress. Euphytica 216:63. 10.1007/s10681-020-02599-6

Bagale S, Enesi RO, Gorim LY (2025) An overview of root traits and ideotypes for improving crop productivity and addressing agronomic challenges. Rhizosphere 34:101105. 10.1016/j.rhisph.2025.101105

Bahadur A, Singh PM, Rai N, et al (2024) Grafting in vegetables to improve abiotic stress tolerance, yield and quality. J Hortic Sci Biotechnol 99:385–403. 10.1080/14620316.2023.2299009

Barzegar T, Heidaryan N, Lotfi H, Ghahremani Z (2018) Yield, fruit quality and physiological responses of melon cv. Khatooni under deficit irrigation. Adv Hortic Sci 32:451–458

Bradbury PJ, Zhang Z, Kroon DE, et al (2007) TASSEL: software for association mapping of complex traits in diverse samples. Bioinformatics 23:2633–2635. 10.1093/bioinformatics/btm308

Broman KW, Wu H, Sen Ś, Churchill GA (2003) R/qtl: QTL mapping in experimental crosses. Bioinformatics 19:889–890. 10.1093/bioinformatics/btg112

Buckler ES, Holland JB, Bradbury PJ, et al (2009) The genetic architecture of maize flowering time. Science 325:714–718. 10.1126/science.1174276

Courtois B, Ahmadi N, Khowaja F, et al (2009) Rice root genetic architecture: meta-analysis from a drought QTL database. Rice 2:115–128. 10.1007/s12284-009-9028-9

Dafna A, Halperin I, Oren E, et al (2021) Underground heterosis for yield improvement in melon. J Exp Bot 72:6205–6218. 10.1093/jxb/erab219

Danin-Poleg Y, Tadmor Y, Tzuri G, et al (2002) Construction of a genetic map of melon with molecular markers and horticultural traits, and localization of genes associated with ZYMV resistance. Euphytica 125:373–384. 10.1023/A:1016021926815

Davis M, Stone A, Selman L, et al (2024) Grafting onto tomato rootstocks improves outcomes for dry-farmed tomato. HortTechnology 34:313–321. 10.21273/HORTTECH05412-24

Duvick DN (2001) Biotechnology in the 1930s: the development of hybrid maize. Nat Rev Genet 2:69–74. 10.1038/35047587

East EM (1908) Inbreeding in corn. Rep Conn Agric Exp Stn 1907:419–428

Edelstein M, Burger Y, Horev C, et al (2004) Assessing the effect of genetic and anatomic variation of Cucurbita rootstocks on vigour, survival and yield of grafted melons. J Hortic Sci Biotechnol 79:370–374. 10.1080/14620316.2004.11511775

Eshed Y, Zamir D (1996) Less-than-additive epistatic interactions of quantitative trait loci in tomato. Genet 143:1807–1817

Estañ MT, Villalta I, Bolarín MC, et al (2009) Identification of fruit yield loci controlling the salt tolerance conferred by solanum rootstocks. Theor Appl Genet 118:305–312. 10.1007/s00122-008-0900-6

FAO, WHO, IFAD, et al (2025) The state of food security and nutrition in the world 2025: addressing high food price inflation for food security and nutrition. Food & Agriculture Org.

Fita A, Picó B, Monforte AJ, Nuez F (2008) Genetics of root system architecture using near isogenic lines of melon. J Am Soc Hortic Sci 133:448–458. 10.21273/JASHS.133.3.448

Fitter A (2002) Characteristics and functions of root systems. In: Plant Roots, 3rd edn. CRC Press, pp 49–78

Flores-León A, García-Martínez S, González V, et al (2021) Grafting snake melon [Cucumis melo L. subsp. melo var. flexuosus (L.) naudin] in organic farming: effects on agronomic performance resistance to pathogens sugar acid and VOC profiles and consumer acceptance. Front Plant Sci 12:. 10.3389/fpls.2021.613845

Galpaz N, Gonda I, Shem-Tov D, et al (2018) Deciphering genetic factors that determine melon fruit-quality traits using RNA-Seq-based high-resolution QTL and eQTL mapping. The Plant journal : for cell and molecular biology 94:169–191. 10.1111/tpj.13838

Gamuyao R, Chin JH, Pariasca-Tanaka J, et al (2012) The protein kinase Pstol1 from traditional rice confers tolerance of phosphorus deficiency. Nature 488:535–539. 10.1038/nature11346

Garcia AAF, Wang S, Melchinger AE, Zeng Z-B (2008) Quantitative trait loci mapping and the genetic basis of heterosis in maize and rice. Genetics 180:1707–1724. 10.1534/genetics.107.082867

Garcia-Mas J, Benjak A, Sanseverino W, et al (2012) The genome of melon (Cucumis melo L.). Proc Natl Acad Sci U S A 109:11872–11877. 10.1073/pnas.1205415109

Godfray HCJ, Beddington JR, Crute IR, et al (2010) Food security: the challenge of feeding 9 billion people. Science 327:812–818. 10.1126/science.1185383

Gomez-Roldan V, Fermas S, Brewer PB, et al (2008) Strigolactone inhibition of shoot branching. Nature 455:189–194. 10.1038/nature07271

Gong T, Brecht JK, Koch KE, et al (2022) A systematic assessment of how rootstock growth characteristics impact grafted tomato plant biomass, resource partitioning, yield, and fruit mineral composition. Front Plant Sci 13:. 10.3389/fpls.2022.948656

Gonzalo MJ, Díaz A, Dhillon NPS, et al (2019) Re-evaluation of the role of Indian germplasm as center of melon diversification based on genotyping-by-sequencing analysis. BMC Genomics 20:448. 10.1186/s12864-019-5784-0

Guo M, Rupe MA, Wei J, et al (2014) Maize ARGOS1 (ZAR1) transgenic alleles increase hybrid maize yield. J Exp Bot 65:249–260. 10.1093/jxb/ert370

Gur A, Semel Y, Osorio S, et al (2011) Yield quantitative trait loci from wild tomato are predominately expressed by the shoot. Theor Appl Genet 122:405–420. 10.1007/s00122-010-1456-9

Gur A, Zamir D (2015) Mendelizing all components of a pyramid of three yield QTL in tomato. Front Plant Sci 6:1–13. 10.3389/fpls.2015.01096

Harel-Beja R, Tzuri G, Portnoy V, et al (2010) A genetic map of melon highly enriched with fruit quality QTLs and EST markers, including sugar and carotenoid metabolism genes. Theor Appl Genet 121:511–533. 10.1007/s00122-010-1327-4

Hodge A (2004) The plastic plant: root responses to heterogeneous supplies of nutrients. New Phytol 162:9–24. 10.1111/j.1469-8137.2004.01015.x

Hua J, Xing Y, Wu W, et al (2003) Single-locus heterotic effects and dominance by dominance interactions can adequately explain the genetic basis of heterosis in an elite rice hybrid. Proc Natl Acad Sci U S A 100:2574–2579. 10.1073/pnas.0437907100

Huang X, Yang S, Gong J, et al (2015) Genomic analysis of hybrid rice varieties reveals numerous superior alleles that contribute to heterosis. Nat Commun 6:6258. 10.1038/ncomms7258

Huntenburg K, Puértolas J, de Ollas C, Dodd IC (2022) Bi-directional, long-distance hormonal signalling between roots and shoots of soil water availability. Physiol Plant 174:e13697. 10.1111/ppl.13697

Ingram TW, Sharpe S, Trandel M, et al (2022) Vigorous rootstocks improve yields and increase fruit sizes in grafted fresh market tomatoes. Front Hortic 1:. 10.3389/fhort.2022.1091342

Kabas A, Uluisik S, Ustun H, et al (2025) Solanum lycopersicoides introgression lines used as rootstocks uncover QTLs affecting tomato morphological and fruit quality traits. Horticulturae 11:. 10.3390/horticulturae11111364

Koevoets IT, Venema JH, Elzenga JTM, Testerink C (2016) roots withstanding their environment: exploiting root system architecture responses to abiotic stress to improve crop tolerance. Front Plant Sci 7:. 10.3389/fpls.2016.01335

Krieger U, Lippman ZB, Zamir D (2010) The flowering gene SINGLE FLOWER TRUSS drives heterosis for yield in tomato. Nat Genet 42:459–463. 10.1038/ng.550

Lippman ZB, Zamir D (2007) Heterosis: revisiting the magic. Trends Genet 23:60–66. 10.1016/j.tig.2006.12.006

Liu J, Li M, Zhang Q, et al (2020) Exploring the molecular basis of heterosis for plant breeding. J Integr Plant Biol 62:287–298. 10.1111/jipb.12804

Lynch JP (2019) Root phenotypes for improved nutrient capture: an underexploited opportunity for global agriculture. New Phytol 223:548–564. 10.1111/nph.15738

Lynch JP (2013) Steep, cheap and deep: an ideotype to optimize water and N acquisition by maize root systems. Ann Bot 112:347–357. 10.1093/aob/mcs293

Marcelis LFM, Heuvelink E, Baan Hofman-Eijer LR, et al (2004) Flower and fruit abortion in sweet pepper in relation to source and sink strength. J Exp Bot 55:2261–2268. 10.1093/jxb/erh245

McMullen MD, Kresovich S, Villeda HS, et al (2009) Genetic properties of the maize nested association mapping population. Science 325:737–740. 10.1126/science.1174320

Ndlovu ME, Soundy P, Klerk JJD, et al (2025) Yield and quality response of indeterminate tomatoes to combined growing methods and rootstock cultivars. Horticulturae 11:. 10.3390/horticulturae11070758

Oren E, Dafna A, Tzuri G, et al (2022) Pan-genome and multi-parental framework for high-resolution trait dissection in melon (Cucumis melo). Plant J 112:1525–1542. 10.1111/tpj.16021

Oren E, Tzuri G, Dafna A, et al (2020) High-density NGS-based map construction and genetic dissection of fruit shape and rind netting in Cucumis melo. Theor Appl Genet 133:1927–1945. 10.1007/s00122-020-03567-3

Osakabe Y, Yamaguchi-Shinozaki K, Shinozaki K, Tran L-SP (2014) ABA control of plant macroelement membrane transport systems in response to water deficit and high salinity. New Phytol 202:35–49. 10.1111/nph.12613

Paez-Garcia A, Motes CM, Scheible W-R, et al (2015) Root traits and phenotyping strategies for plant improvement. Plants 4:334–355. 10.3390/plants4020334

Paril J, Reif J, Fournier-Level A, Pourkheirandish M (2024) Heterosis in crop improvement. Plant J 117:23–32. 10.1111/tpj.16488

Parry MAJ, Hawkesford MJ (2010) Food security: increasing yield and improving resource use efficiency. Proc Nutr Soc 69:592–600. 10.1017/S0029665110003836

Pitrat M (2017) Melon genetic resources: phenotypic diversity and horticultural taxonomy. In: Grumet R, Katzir N, Garcia-Mas J (eds) Genetics and Genomics of Cucurbitaceae. Springer International Publishing, Cham, pp 25–60

Sakakibara H (2006) CYTOKININS: activity, biosynthesis, and translocation. Annu Rev Plant Biol 57:431–449. 10.1146/annurev.arplant.57.032905.105231

Satasiya P, Patel S, Patel R, et al (2024) Meta-analysis of identified genomic regions and candidate genes underlying salinity tolerance in rice (Oryza sativa L.). Sci Rep 14:5730. 10.1038/s41598-024-54764-9

Schnable PS, Springer NM (2013) Progress toward understanding heterosis in crop plants. Annu Rev Plant Biol 64:71–88. 10.1146/annurev-arplant-042110-103827

Semel Y, Nissenbaum J, Menda N, et al (2006) Overdominant quantitative trait loci for yield and fitness in tomato. Proc Natl Acad Sci U S A 103:12981–12986. 10.1073/pnas.0604635103

Shull GH (1908) The composition of a field of maize. J Hered os-4:296–301. 10.1093/jhered/os-4.1.296

Smith S, De Smet I (2012) Root system architecture: insights from Arabidopsis and cereal crops. Phil Trans R Soc B 367:1441–1452. 10.1098/rstb.2011.0234

Tilman D, Balzer C, Hill J, Befort BL (2011) Global food demand and the sustainable intensification of agriculture. Proc Natl Acad Sci U S A 108:20260–20264. 10.1073/pnas.1116437108

Tracy SR, Nagel KA, Postma JA, et al (2020) Crop improvement from phenotyping roots: highlights reveal expanding opportunities. Trends Plant Sci 25:105–118. 10.1016/j.tplants.2019.10.015

Tzuri G, Dafna A, Itzhaki B, et al (2025) Meta genetic analysis of melon sweetness. Theor Appl Genet 138: 68. 10.1007/s00122-025-04863-6

Uga Y, Sugimoto K, Ogawa S, et al (2013) Control of root system architecture by DEEPER ROOTING 1 increases rice yield under drought conditions. Nat Genet 45:1097–1102. 10.1038/ng.2725

Wang H, Qi M, Cutler AJ (1993) A simple method of preparing plant samples for PCR. Nucleic Acids Res 21:4153. 10.1093/nar/21.17.4153

Wasson AP, Rebetzke GJ, Kirkegaard JA, et al (2014) Soil coring at multiple field environments can directly quantify variation in deep root traits to select wheat genotypes for breeding. J Exp Bot 65:6231–6249. 10.1093/jxb/eru250

Xiao J, Li J, Yuan L, Tanksley SD (1995) Dominance is the major genetic basis of heterosis in rice as revealed by QTL analysis using molecular markers. Genetics 140:745–754. 10.1093/genetics/140.2.745

Xiao Y, Jiang S, Cheng Q, et al (2021) The genetic mechanism of heterosis utilization in maize improvement. Genome Biol 22:148. 10.1186/s13059-021-02370-7

Zhao G, Lian Q, Zhang Z, et al (2019) A comprehensive genome variation map of melon identifies multiple domestication events and loci influencing agronomic traits. Nat Genet 51:1607–1615. 10.1038/s41588-019-0522-8

Zhu J, Kaeppler SM, Lynch JP (2005) Mapping of QTLs for lateral root branching and length in maize (Zea mays L.) under differential phosphorus supply. Theor Appl Genet 111:688–695. 10.1007/s00122-005-2051-3

