## Supplemental Figures S1-2 for "Genetic dissection of root-mediated yield heterosis in melon (*Cucumis melo*)"

Figure S1

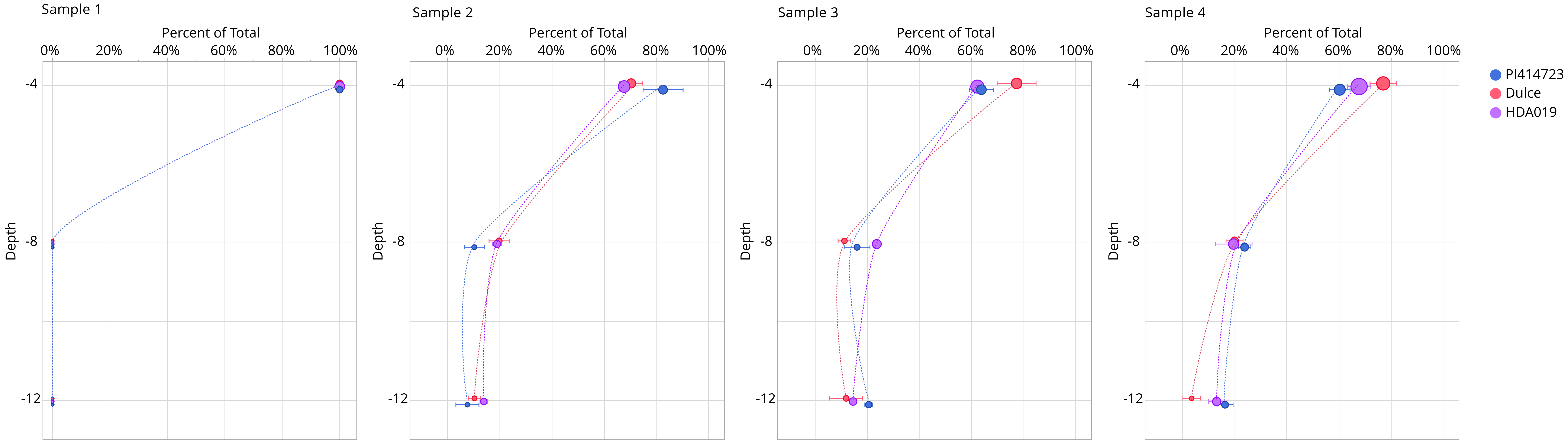
Sample 1
Sample 2
Sample 3
Sample 4
Percent of Total
0%
20%
40%
60%
80%
100%
Depth
-4
-8
-12
PI414723
Dulce
HDA019

Figure S2

Fruits of PI414723 Parental line

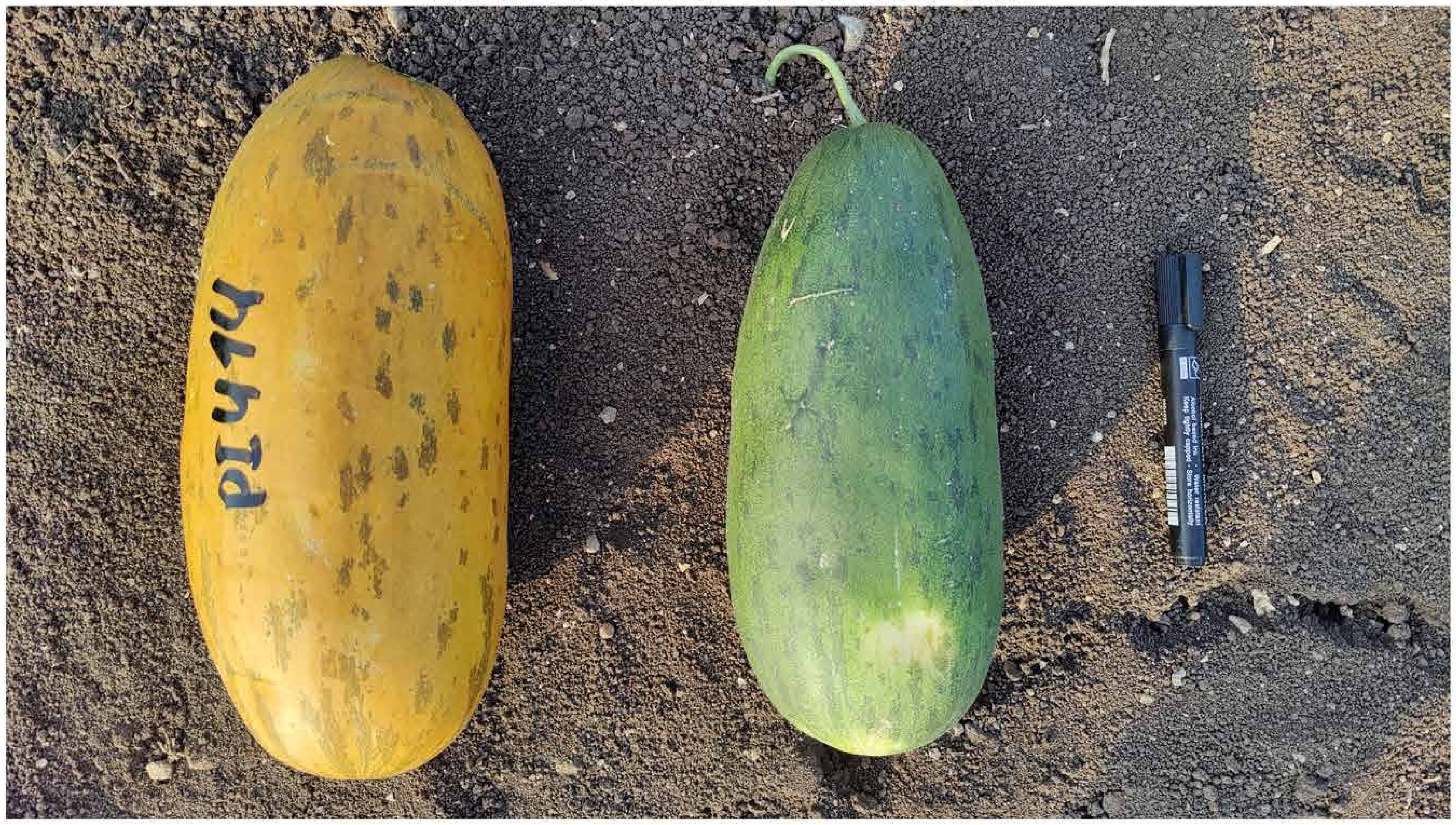


Fruits of two BC3F3 lines with favorable (DUL) allele in QTL10.3

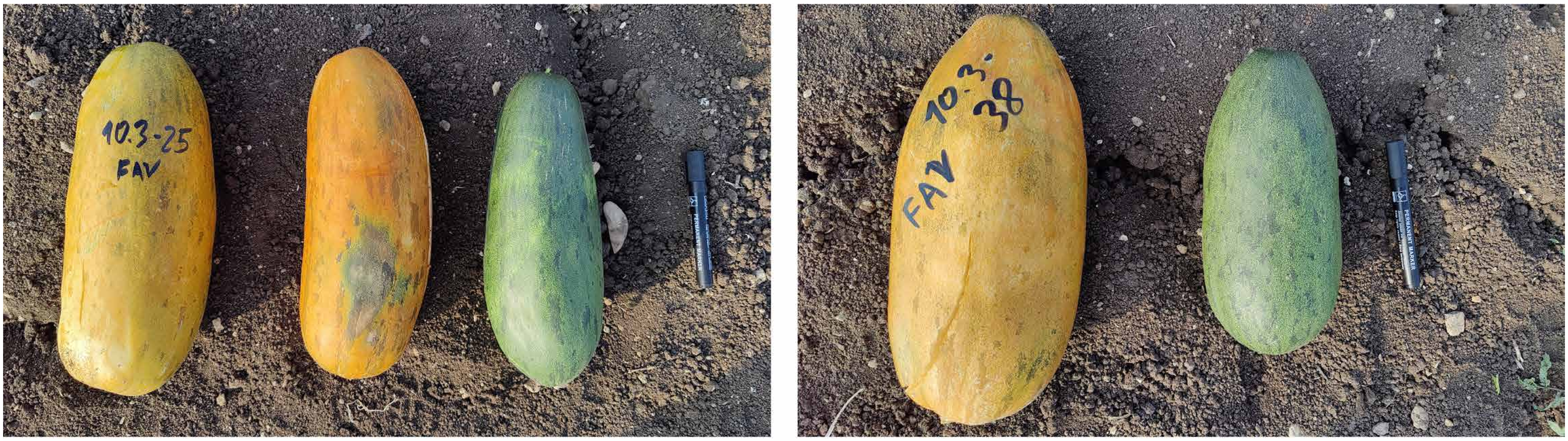


Fruits of BC3F3 line with favorable (DUL) allele in QTL2.4

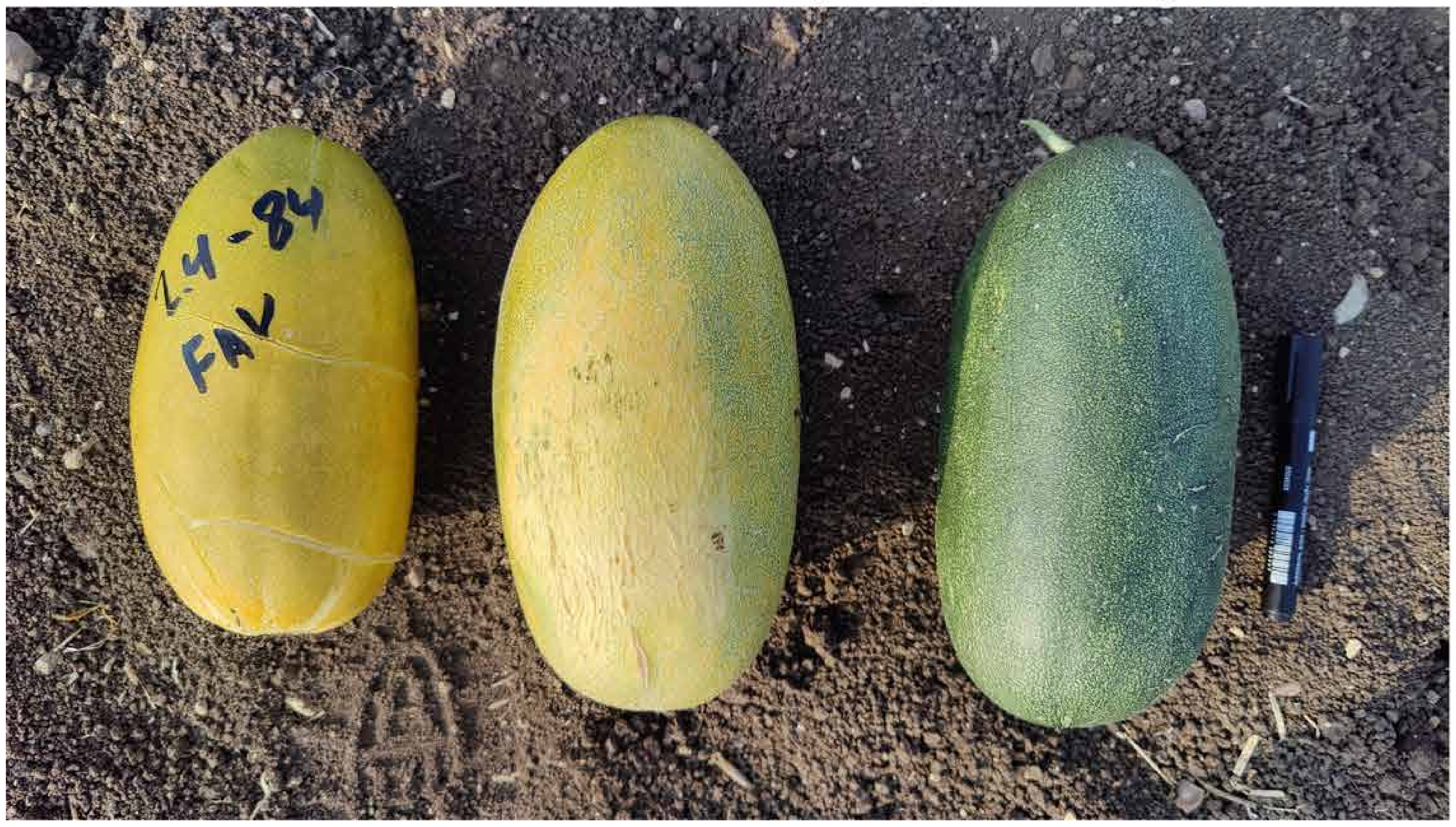


Fruits of BC3F3 line with non-favorable (PI414) allele in QTL2.4

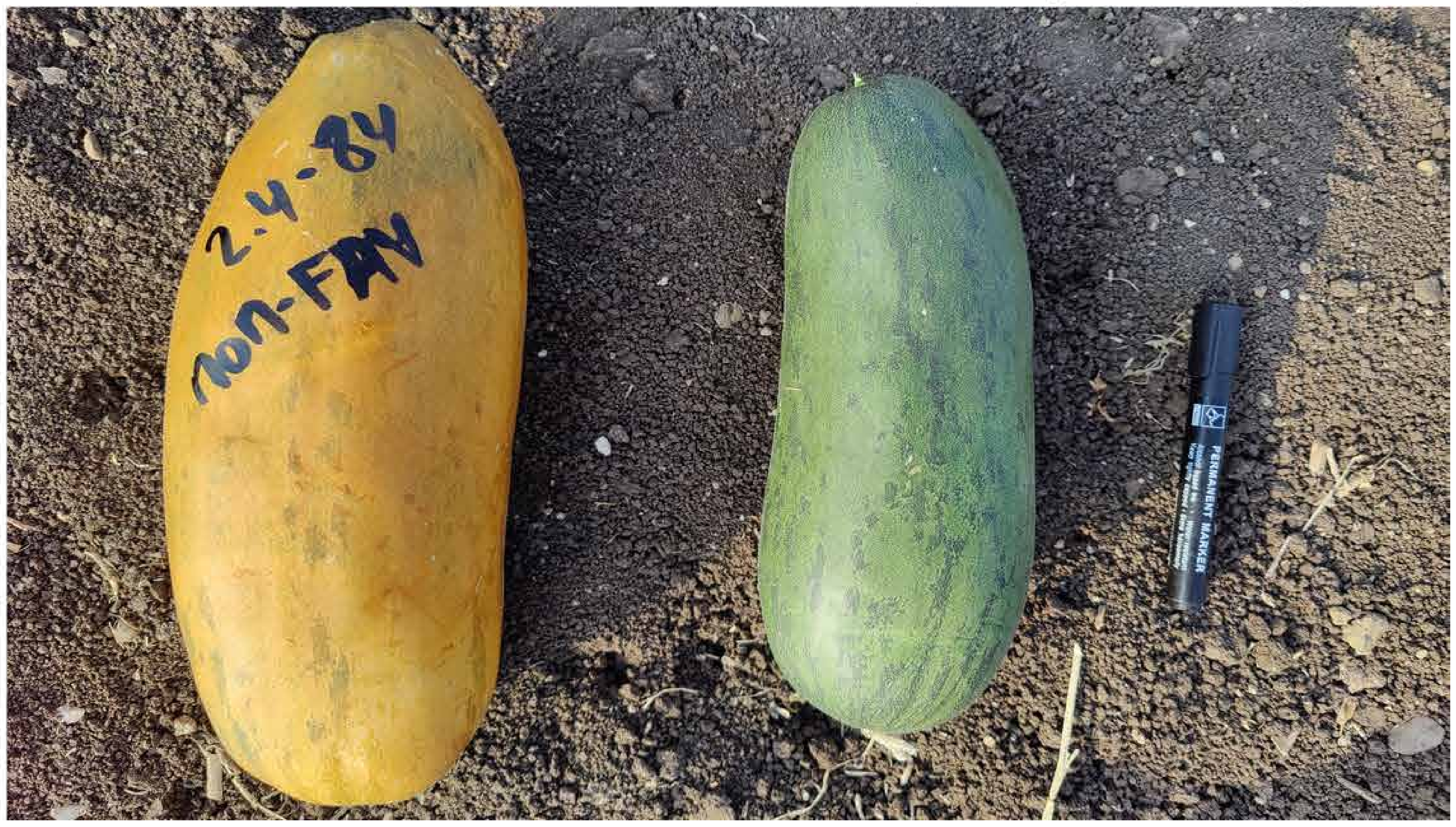
